# Rescue of lysosomal acid lipase deficiency in mice by rAAV8 liver gene transfer

**DOI:** 10.1101/2024.04.26.591270

**Authors:** Marine Laurent, Rim Harb, Christine Jenny, Julie Oustelandt, Simon Jimenez, Jeremie Cosette, Francesca Landini, Aristide Ferrante, Guillaume Corre, Nemanja Vujic, Claudia Piccoli, Anais Brassier, Laetitia Van Wittenberghe, Giuseppe Ronzitti, Dagmar Kratky, Consiglia Pacelli, Mario Amendola

## Abstract

Lysosomal acid lipase deficiency (LAL-D) is an autosomal recessive disorder caused by mutations in the *LIPA* gene, which results in lipid accumulation leading to multi-organ failure. If left untreated, the severe form of LAL-D results in premature death within the first year of life due to failure to thrive and hepatic insufficiency. Enzyme replacement therapy is the only available supportive treatment consisting in weekly systemic injections of recombinant LAL protein. Here, we characterized a novel *Lipa^-/-^* mouse model and developed a curative gene therapy treatment based on the *in vivo* administration of recombinant (r)AAV8 vector encoding the human *LIPA* transgene under the control of a hepatocyte-specific promoter. We defined the minimal rAAV8 dose required to rescue disease lethality and to correct cholesterol and triglyceride accumulation in multiple organs and blood. Finally, using liver transcriptomic and biochemical analysis, we showed mitochondrial impairment in *Lipa^-/-^* mice and its recovery by gene therapy. Overall, our *in vivo* gene therapy strategy achieves a stable long-term LAL expression sufficient to correct the disease phenotype in the *Lipa^-/-^*mouse model and offers a new therapeutic option for LAL-D patients.

**One Sentence Summary:** We’ve developed a liver-targeted gene therapy using recombinant AAV8 to effectively cure Lysosomal acid lipase deficiency by correcting lipid accumulation and by normalizing gene expression pattern and mitochondrial function in *Lipa^-/-^* mouse model.

## INTRODUCTION

Lysosomal acid lipase deficiency (LAL-D) is a rare autosomal recessive lysosomal storage disorder caused by mutations in the *LIPA* gene. In healthy lysosomes, LAL is the sole enzyme responsible for the hydrolysis of cholesterol esters and triglycerides into free cholesterol and fatty acids respectively, necessary for energy production, membrane formation and biosynthesis of cholesterol-derived compounds. In LAL-D patients, cholesterol esters and triglycerides accumulate in tissues and organs, especially in liver, spleen, and small intestine, resulting in multi-organ failure. As a consequence, LAL-D patients suffer from hepatosplenomegaly, elevated circulating alanine (ALT) and aspartate (AST) transaminases due to liver damage and hyperlipidemia with elevated low-density (LDL) and reduced high-density lipoprotein (HDL). Unlike other lysosomal storage disorders, LAL-D does not obviously affect the central nervous system (*1*, *2*). The severity of the disorder depends on the residual amount of functional LAL enzyme. Two forms of the disease have been described: infantile-onset, formerly known as Wolman disease (WD, OMIM 620151; prevalence ∼0.325–1.11 cases per million births), with <2% of remaining functional LAL (*3*, *4*); childhood/adult-onset LAL-D, formerly known as cholesteryl ester storage disease (CESD, OMIM 278000; prevalence 3.13–4.86 cases per million births), with 2-12% of remaining functional LAL. WD patients have difficulty to thrive, intestinal malabsorption, adrenal gland calcification and experience premature death in the first year after birth without treatment. In contrast, CESD patients present milder symptoms, eventually with premature atherosclerosis and cardiac complications that can result in death in childhood or adolescence if left untreated (*4*).

Several therapeutic strategies have been attempted, including hematopoietic stem cells (HSC) and liver transplantation from healthy donors, but they were not effective in preventing disease progression and are limited by high procedure morbidity and HLA matched donor availability (*5–7*). Nowadays, enzyme replacement therapy (ERT) with weekly intravenous administrations of recombinant LAL enzyme Sebelipase (Kanuma®, Alexion Pharmaceuticals, Inc.), is the standard care for LAL-D. This non-curative treatment improves patients’ lifespan and major clinical symptoms. However, it is associated with the development of antibodies against the recombinant enzyme in a fraction of treated patients, it remains very costly and its long-term efficacy is still under evaluation (*8*, *9*).

*In vivo* gene therapy (GT) based on the delivery of a functional copy of the *LIPA* transgene represents an appealing curative treatment to continuously express therapeutic LAL enzyme. A proof-of-principle was previously obtained by intravenous injection of first-generation adenoviral vectors (AdV) encoding for human *LIPA* cDNA under the control of the viral CMV promoter into *Lipa^-/-^* mice (*10*). Transduced hepatocytes expressed and secreted human LAL into the circulation to cross-correct all affected tissues, but the effect was transient and a strong AdV immune response was observed.

To address these constraints, we assessed a liver-based *in vivo* GT for LAL-D using recombinant single-stranded adeno associated viral vector (rAAV) 8 encoding for *LIPA* cDNA under the control of liver promoter. As animal model, we described a new *Lipa^-/-^* mouse characterised by reduced growth, body weight and lifespan (6-7 months) compared to their WT littermates. Upon rAAV8 injection, we established the minimal vector dose required to achieve full correction of hepatosplenomegaly, haematological parameters, dyslipidaemia, and transcriptional deregulation, resulting in normal growth and lifespan of *Lipa^-/-^*mice.

## RESULTS

### rAAV8 injection improves body weight, haematological and plasma parameters at low dose

As disease model, we used a *Lipa^-/-^* mouse model generated by deleting exon 4 of *Lipa* gene, with virtually no residual LAL expression or activity (Fig. S1) (*11*). *Lipa^-/-^* mice failed to thrive and remained smaller compared to WT mice. A recombinant single-stranded adeno associated viral vector (rAAV) 8 was used to deliver the human *LIPA* cDNA under the control of the hepatocyte-specific human α1-antitrypsin (hAAT) promoter (rAAV8.hAAT.LIPA). This rAAV8 construct was intravenously administered into 10–12-week-old male *Lipa^-/-^* mice at 4 different doses: 5×10^11^ viral genome per kilogram (vg/kg) (low dose (LD) 1), 1×10^12^ vg/kg (LD2), 3×10^12^ vg/kg (middle dose; MD), 1×10^13^ vg/kg (high dose; HD). Body weight, LAL activity, haematological parameters, hepatic transaminase (ALT, AST; liver damage markers) and high-density lipoprotein (HDL)-cholesterol concentration were monitored every other week for 12-weeks post-administration (Fig. 1.A). Soon after injection, we observed that treated mice gained body weight according to the injected rAAV8 dose compared to *Lipa^-/-^* mice (Fig. 1.B). Since our approach depends on the hepatic expression and secretion of therapeutic LAL enzyme in the circulation for systemic cross-correction, we measured the LAL activity in plasma. As expected, LAL activity was significantly increased 2-weeks post-injection and positively correlated with the injected rAAV8 dose, reaching up to 4-fold higher activity in circulation compared to WT mice (Fig. 1.C). Noteworthy, LD1 did not result in any increase of LAL activity, while LD2 reached a plateau at 8-weeks post injection. Some LAL activity was still observed in *Lipa^-/-^* mice, probably due to unspecific signals from the high abundance of other lipases in plasma.

**Fig. 1.**
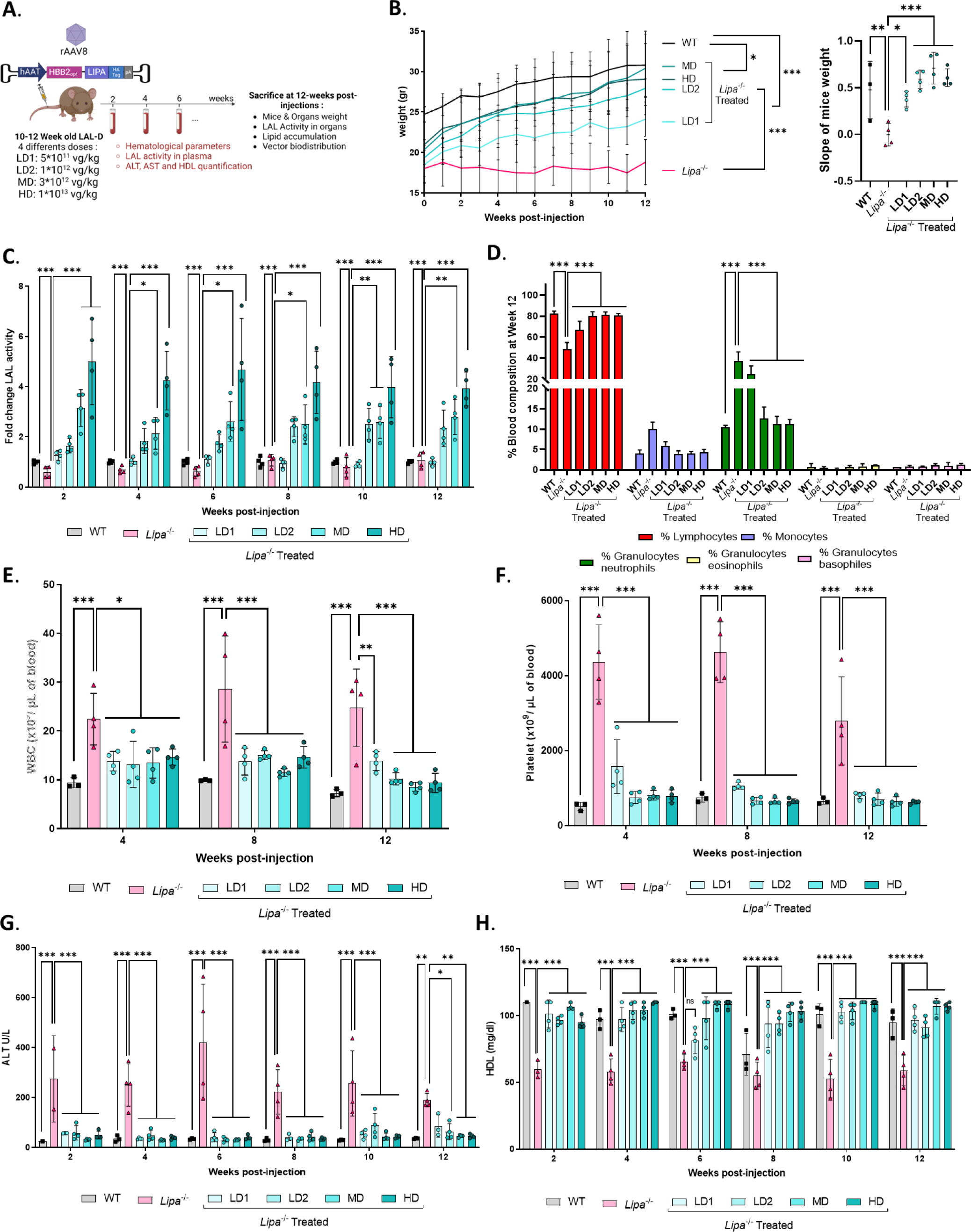
Evaluation of different rAAV8 doses in *Lipa^-/-^* mice. **(A)** Experimental outline created by BioRender.com. 10–12-week-old mice were injected with the indicated rAAV8 doses and disease progression was monitored for 12-weeks post-injection. LD, low dose; MD, middle dose; HD, high dose; vg, vector genome. **(B)** Left panel: Mice weight (gr: grams) was recorded every week. Right panel: Linear slopes of mice weight. Statistical significance was calculated using one-way ANOVA with Kruskal-Wallis Test (*p<0.033; **p<0.002; *** p<0.0002). **(C)** Fold change of LAL activity over WT mice measured in plasma samples every 2-weeks post-injection. **(D)** Blood cell populations in blood samples collected at 12-weeks post-injection. **(E** to **F) (E)** White blood cell (WBC) and **(F)** platelet count in blood samples collected every 4-weeks post-injection. **(G** to **H)** Quantification of **(G)** ALT (U/L, unit per liter) and **(H)** HDL in plasma samples collected every 2-weeks post-injection. Bars indicate mean ± SD (n = 3-4) and statistical significance was calculated using two-way ANOVA with Tukey’s test (*p<0.033; **p<0.002; *** p<0.0002).

Next, we monitored haematological parameters at 4-, 8-and 12-weeks post injection. In *Lipa^-/-^* mice, we observed a shift in blood cell composition (Fig. 1.D, Fig. S2.A), with a significant increase in white blood cells (WBCs) (Fig. 1.E), as previously described (*12*, *13*). This inflammatory profile was confirmed by thrombocytosis (Fig. 1.F) with larger platelets (Fig. S2.B) and microcytic anemia (Fig. S2.C-E). In addition, we observed an increase in AST and ALT concentrations (Fig. 1H, Fig. S2.F), indicating liver damage, and a decrease in HDL (Fig. 1G), a major risk factor for cardiovascular disease. Interestingly, all parameters were fully corrected within 2-weeks post-injection with all rAAV8 doses, except for LD1, which showed a partial rescue.

### Hepatic transduction restores LAL expression in a dose dependent manner and normalizes hepatosplenomegaly

Twelve-weeks after rAAV8 injection, the mice were sacrificed to evaluate the transduction efficiency, LAL enzymatic activity and hepatosplenomegaly. As expected for the rAAV8 serotype, vector genomes were higher in the liver than in the spleen and jejunum, and they were proportional to the injected rAAV8 dose (Fig. 2.A-B; Fig. S3.A). The same trend was observed for LAL enzymatic activity (Fig. 2.C-D, Fig. S3.B), with the lowest hepatic transduction rate (LD1; 0.5 +/-0.1 copy per diploid genome, mean + SD) resulting in a LAL activity around 35% of WT level (Fig. 2.C). Noteworthy, the LAL activity observed in the spleen and the jejunum could be the result of rAAV8 transduction or cross-correction from hepatocyte-secreted LAL into the circulation. We then assessed the efficacy of LAL in improving the affected organs. Hepatosplenomegaly (Fig. 2.E-G) and liver discoloration (Fig. 2.G) due to increased lipid accumulation in *Lipa^-/-^* mice were significantly decreased with each of the four rAAV8 doses tested (Fig. 2.E-G). Interestingly, the LD1 reduced organs weight, although liver discoloration was only partially corrected at 12-weeks post-administration (Fig. 2.G).

**Fig. 2.**
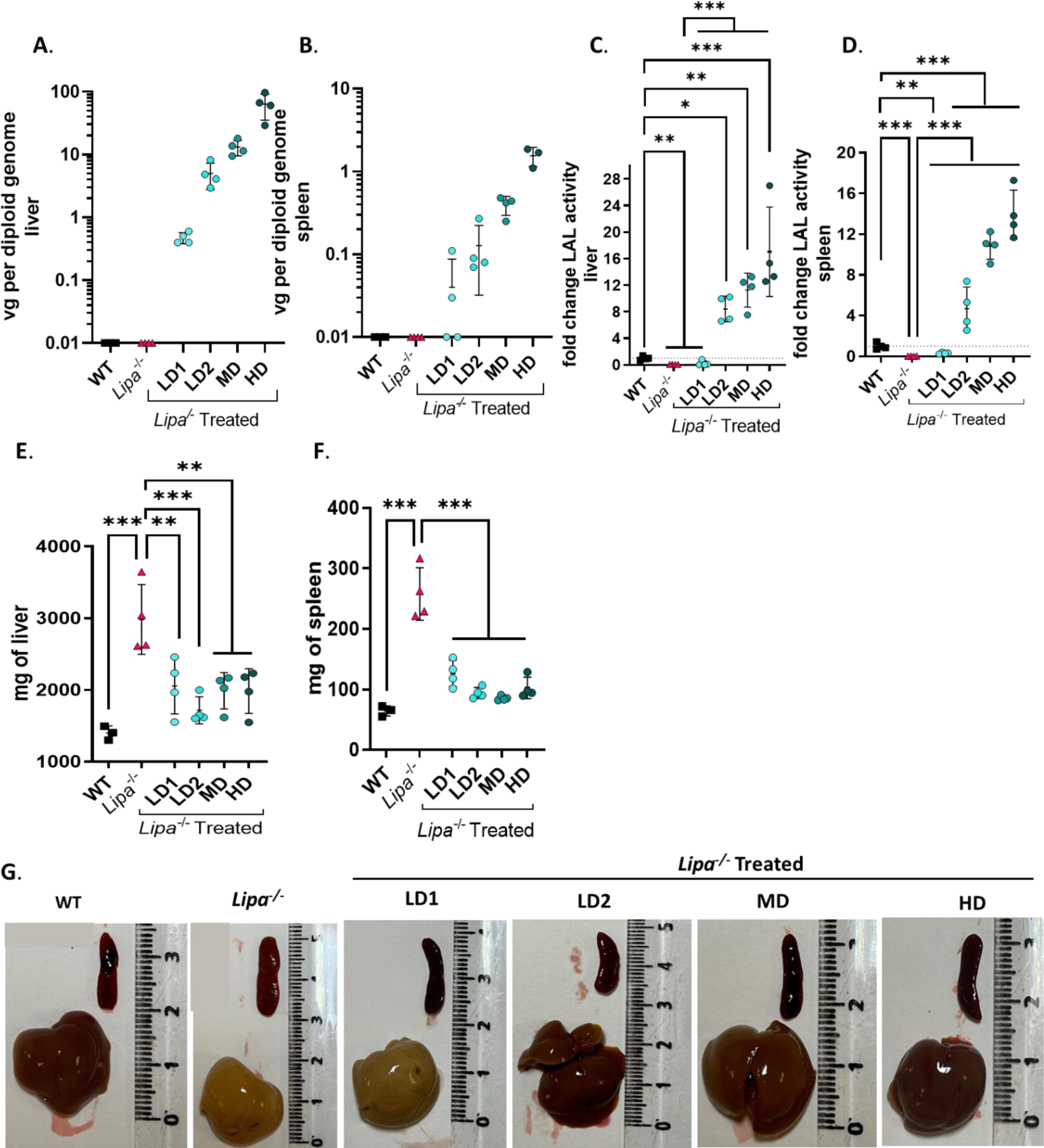
rAAV8 transduction of liver and spleen of *Lipa^-/-^* mice increases LAL activity and corrects hepatosplenomegaly. **(A** *to* **B)** rAAV8 transduction level for different vector doses at 12-weeks post injection **(A)** in liver and **(B)** spleen. vg = vector genome. **(C** *to* **D)** Fold change of LAL activity over WT mice **(C)** in liver and **(D)** spleen for different vector doses. **(E** *to* **F)** Weight of **(E)** liver and **(F)** spleen at 12-weeks post-injection for different vector doses. **(G)** Representative images of liver and spleen at 12-weeks post-injection. Ruler indicates centimetres. Bars indicate mean ± SD (n = 3-4) and statistical significance was calculated using two-way ANOVA with Tukey’s test (*p<0.033; **p<0.002; *** p<0.0002).

Overall, these findings demonstrate that rAAV8 injection efficiently transduced the liver and corrected hepatosplenomegaly.

### LAL overexpression restores lipid homeostasis in liver and spleen

In *Lipa*^-/-^ mice, hepatosplenomegaly is caused by massive accumulation of total cholesterol and triglycerides (Fig. 3.A-D; Fig. S3.C-F). 12-weeks post-treatment, all rAAV8 doses fully reverted cholesterol and triglyceride content in the liver of *Lipa*^-/-^ mice, with the exception of LD1, which reduced cholesterol deposits by more than half (Fig. 3.A; Fig. S3.C). Similar results were obtained for the spleen, although lipid accumulation was less pronounced than in the liver (Fig. 3.C-D; Fig. S3.E-F). In the jejunum, we observed only cholesterol accumulation, which was corrected upon rAAV8 treatment (Fig. S3.G-H).

**Fig. 3.**
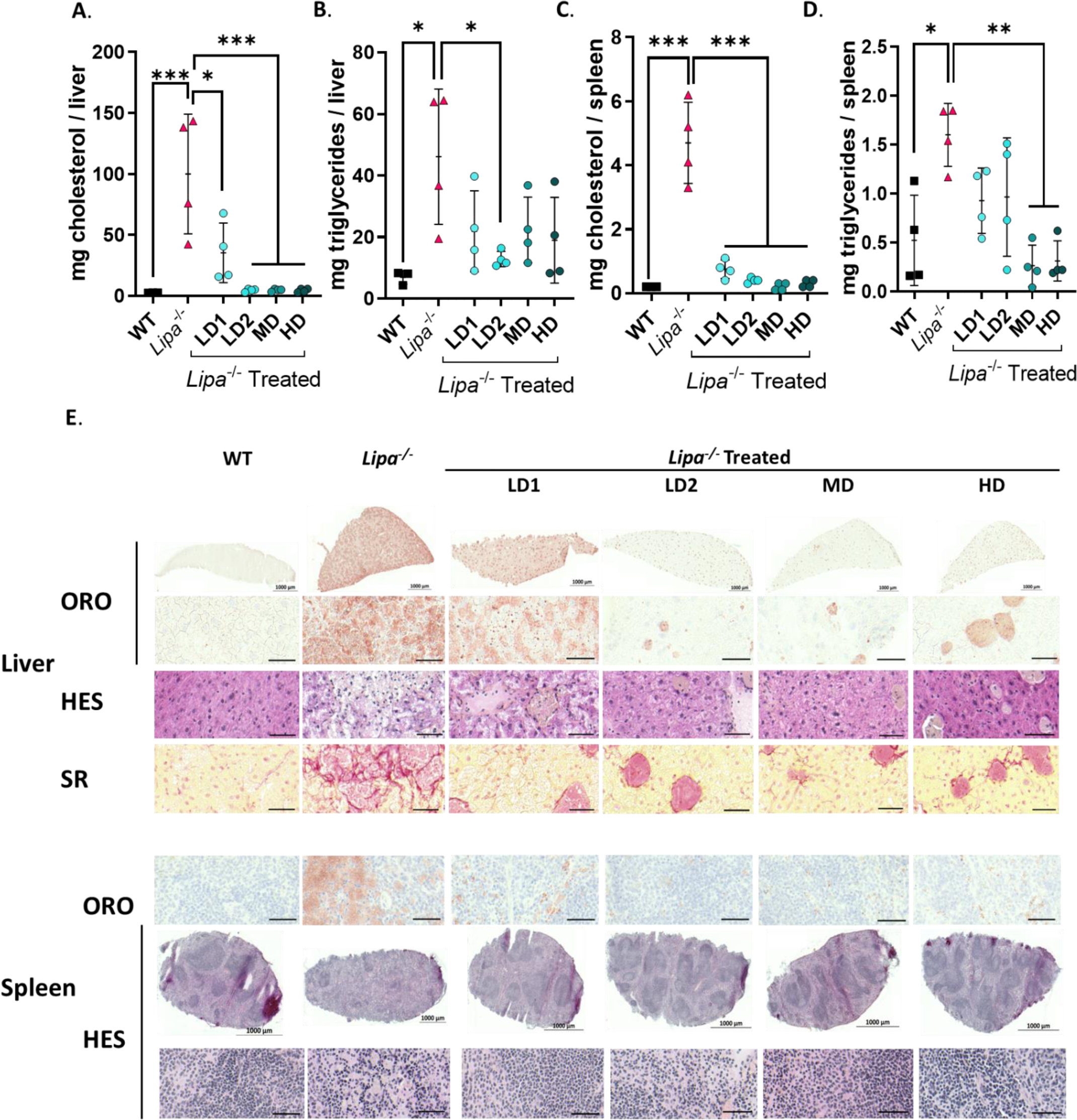
rAAV8 administration corrects lipid accumulation in liver and spleen. **(A** to **D)** Quantification of cholesterol and triglycerides in **(A** to **B)** liver and **(C** to **D)** spleen at 12-weeks post-injection. mg = milligrams. Bars indicate mean ± SD (n=3-4) and statistical significance was calculated using one-way ANOVA with Tukey’s test (*p<0.033; **p<0.002; *** p<0.0001). **(E)** Histological sections: Oil red O (ORO), Haematoxylin Eosin Saffron (HES) and Sirius red (SR) staining of liver (top panel), spleen (bottom panel) at 12-weeks post-injection (scale bar: 50 µm unless differently noted).

We then performed liver and spleen histological analyses to assess architectural alterations. Oil red O (ORO) staining revealed pronounced accumulations of neutral lipids in *Lipa*^-/-^ mice (Fig. 3.E; Fig. S3.I-J), while haematoxylin and eosin staining revealed an increase in connective tissue and a loss of red and white pulps in spleen (Fig. 3.E). Finally, appearance of fibrosis was observed by Sirius red staining (Fig. 3.E; Fig. S3.K). All these pathological signs were mostly reversed 12-weeks after rAAV8 injection, except for LD1 that showed partial efficacy.

### rAAV8 injection results in improved survival rate and disease correction up to 34-weeks post-treatment

Since MD was the lowest vector dose required to achieve correction of *Lipa*^-/-^ mice at 12-weeks post-injection (Fig. 1-3), we evaluated its therapeutic benefits at longer time points. Hence, we biweekly monitored body weight, LAL activity, haematological parameters, transaminases, and HDL-cholesterol for 34-weeks post-injection and collected tissues and organs at 4-, 12- and 34-weeks post-injection.

Our data demonstrated that rAAV8 administration prolonged survival, improved body weight, and normalized growth of *Lipa*^-/-^ mice for at least 34-weeks post-injection (Fig. 4.A-C). In addition, LAL activity in plasma was 2.5-fold increased (Fig. 4.D), which fully corrected blood cell composition (Fig. 4.E and Fig. S4.A), WBC counts (Fig. 4.F), thrombocytosis (Fig. 4.G and Fig. S4.B), and microcytic anemia (Fig. S4.C-E). Finally, LAL expression reduced ALT (Fig. 4.H) and normalized HDL levels (Fig. 4.I). Noteworthy, correction of haematological and serum parameters was already completed after 2-week post-injection and remained stable throughout the entire 34-weeks follow-up period.

**Fig. 4.**
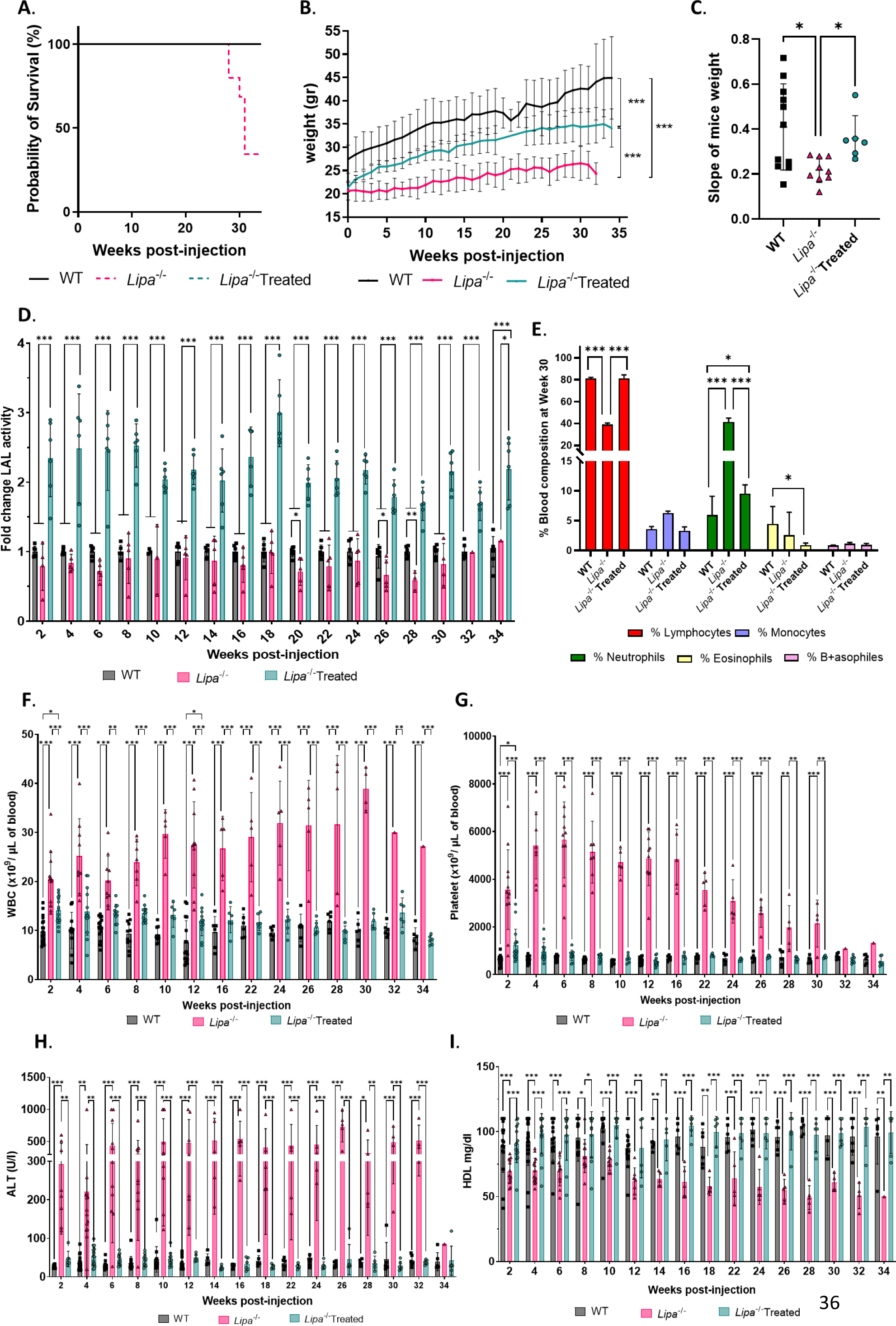
rAAV8 administration increases survival and body weight of *Lipa^-/-^* mice and corrects haematological parameters and dyslipidaemia for 34-weeks post-injection. (A) Percentage of mice survival. **(B)** Mice weight (gr: grams) was recorded every week (n=6-10). **(C)** Linear slopes of mice weight. Bars indicate mean ± SD and statistical significance was calculated using one-way ANOVA with Kruskal-Wallis Test (*p<0.033; **p<0.002; *** p<0.0002). **(D)** Fold change of LAL activity over WT mice measured in plasma samples every 2-weeks post-injection (n=1-20). **(E)** Blood cell populations in blood samples collected at 30-weeks post-injection (n=6-11). **(F** to **G) (F)** White blood cell (WBC) and **(G)** platelet counts in blood samples collected every 2-weeks post-injection (n=1-20). **(H** to **I)** Quantification of **(H)** ALT (U/L, unit per liter) and **(I)** HDL in plasma samples collected every 2-weeks post-injection (n=1-20). Bars indicate mean ± SD and statistical significance was calculated using two-way ANOVA with Tukey’s test (*p<0.033; **p<0.002; *** p<0.0002). ith Tukey’s test (*p<0.033; **p<0.002; *** p<0.0002).

Treated mice were sacrificed at 4-, 12- and 34-weeks post-injection to evaluate rAAV8 transduction efficiency, LAL activity in organs and hepatosplenomegaly over time. As observed previously, we detected higher rAAV8 transduction and LAL expression in the liver compared to the spleen and jejunum at all-time points (Fig. 5.A-D and Fig. S5.A-B). In *Lipa*^-/-^ mice, the liver weight accounted for one-fourth of the total body weight, which was reduced to one-twentieth of the total body weight in rAAV8-treated, as in WT mice (Fig. 5.E, G). The enlargement of both the liver and the spleen was caused by the accumulation of total cholesterol and triglycerides, as quantified biochemically (Fig. 6.A-D and Fig. S5.C-F) and histologically by ORO staining (Fig. 6.E; Fig. S5.G-H). In addition, SR staining revealed architectural alterations and fibrosis in the liver (Fig. 6.E; Fig. S5.I). To a lower extent, lipid accumulation and structural alterations were also observed in the jejunum (Fig. S5.J-L; Fig. S5.M) and thymus (Fig. S6). Noteworthy, organ correction was already visible after 4-weeks post-injection and for at least 34-weeks.

**Fig. 5.**
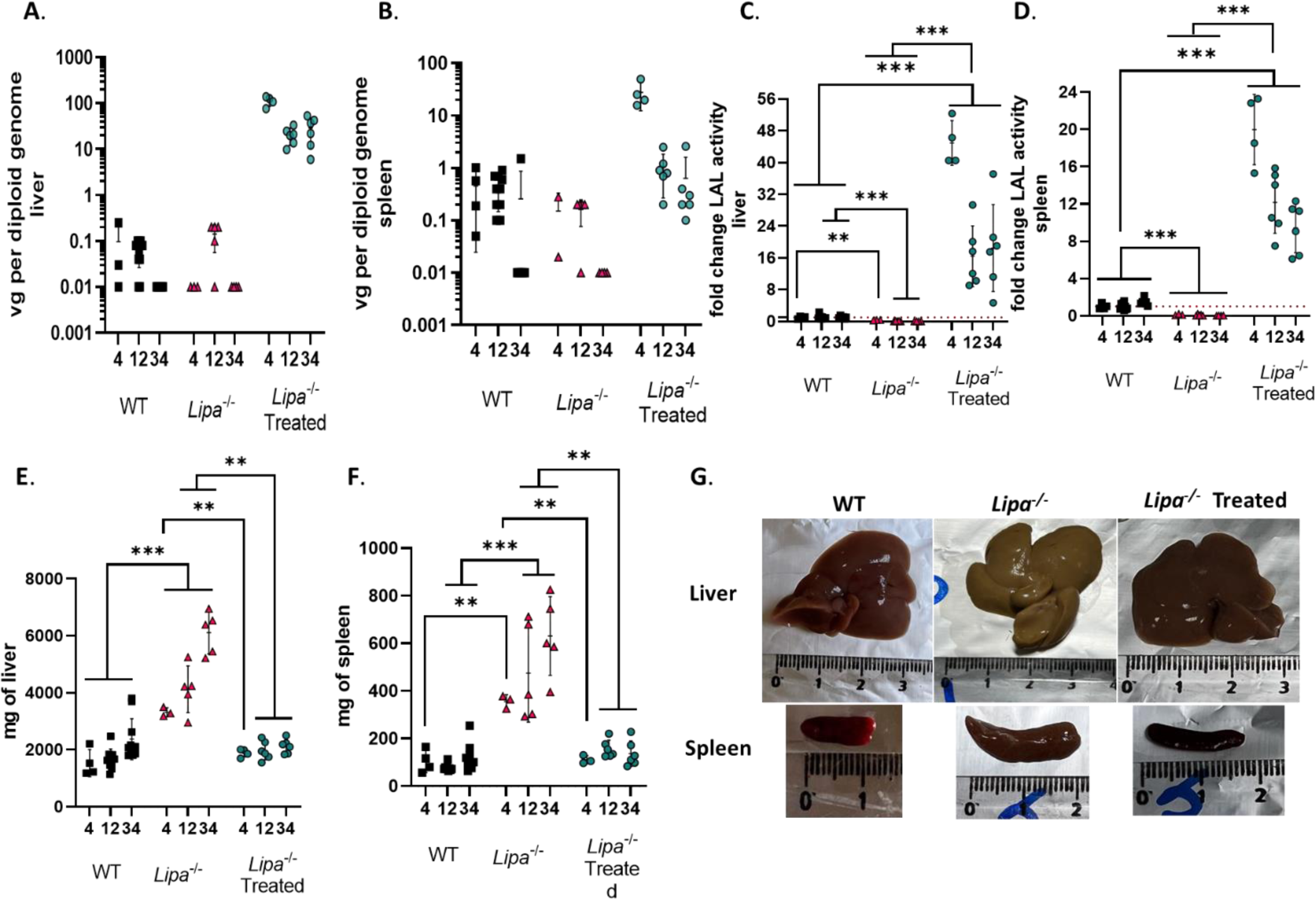
rAAV8 transduction of liver and spleen of *Lipa^-/-^* mice results in long-term increase of LAL activity and correction of hepatosplenomegaly. **(A** to **B)** rAAV8 transduction level in **(A)** liver and **(B)** spleen at indicated time points (weeks) (n=4-10). vg = vector genome. **(C** to **D)** Fold change of LAL activity over WT mice in **(C)** liver and **(D)** spleen at indicated time points (weeks). **(E** to **F)** Weight of **(E)** liver and **(F)** spleen at indicated time points (weeks). **(G)** Representative images of liver and spleen at 12-weeks post-injection. Bars indicate mean ± SD (n = 4-10) and statistical significance was calculated using two-way ANOVA with Tukey’s test (*p<0.033; **p<0.002; *** p<0.0002).

**Fig. 6.**
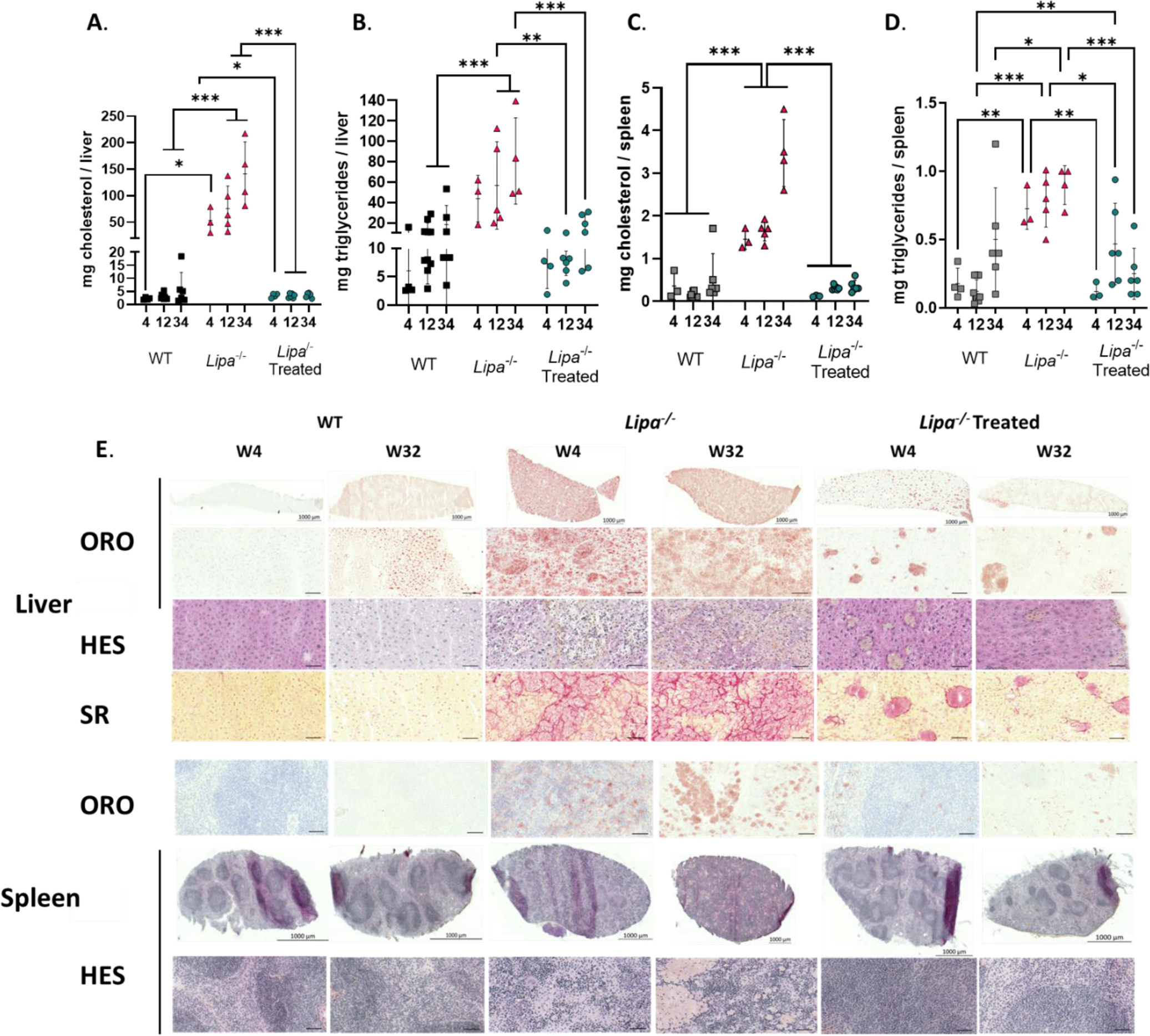
rAAV8 administration corrects lipid accumulation in liver and spleen long-term. **(A** *to* **D)** Quantification of cholesterol and triglycerides in **(A** to **B)** liver and **(C** to **D)** spleen at indicated time points (weeks). Bars indicate mean ± SD (n=4-10). Statistical significance was calculated using two-way ANOVA with Tukey’s test (*p<0.033; **p<0.002; *** p<0.0001). **(E)** Histological sections with: Oil red O (ORO; top row), Haematoxylin Eosin Saffron (HES; middle row) and Sirius red (SR; bottom row) staining of liver (top panel), and spleen (bottom panel) at 4- and 34-weeks (scale: 50 µm).

### rAAV8 injection improves general metabolic, mitochondrial, and immune system dysregulation

Metabolic alterations can have profound and far-reaching effects beyond lipid accumulations. Therefore, to gain a comprehensive understanding, we performed an unbiased transcriptomic analysis of WT, *Lipa*^-/-^ and rAAV8.hAAT.LIPA-treated *Lipa*^-/-^ mouse livers at 12- and 34-weeks post-injection. Principal component analysis (PCA) of gene expression showed that: 1) WT mice at 12- and 34-weeks grouped together, similarly to *Lipa*^-/-^ mice at 12 and 34-week; 2) *Lipa*^-/-^ mice clearly diverged from the other groups at both time points; 3) rAAV8-treated *Lipa*^-/-^ mice diverged from *Lipa*^-/-^ mice and grouped with WT mice at 34-weeks (Fig. 7.A).

**Fig. 7.**
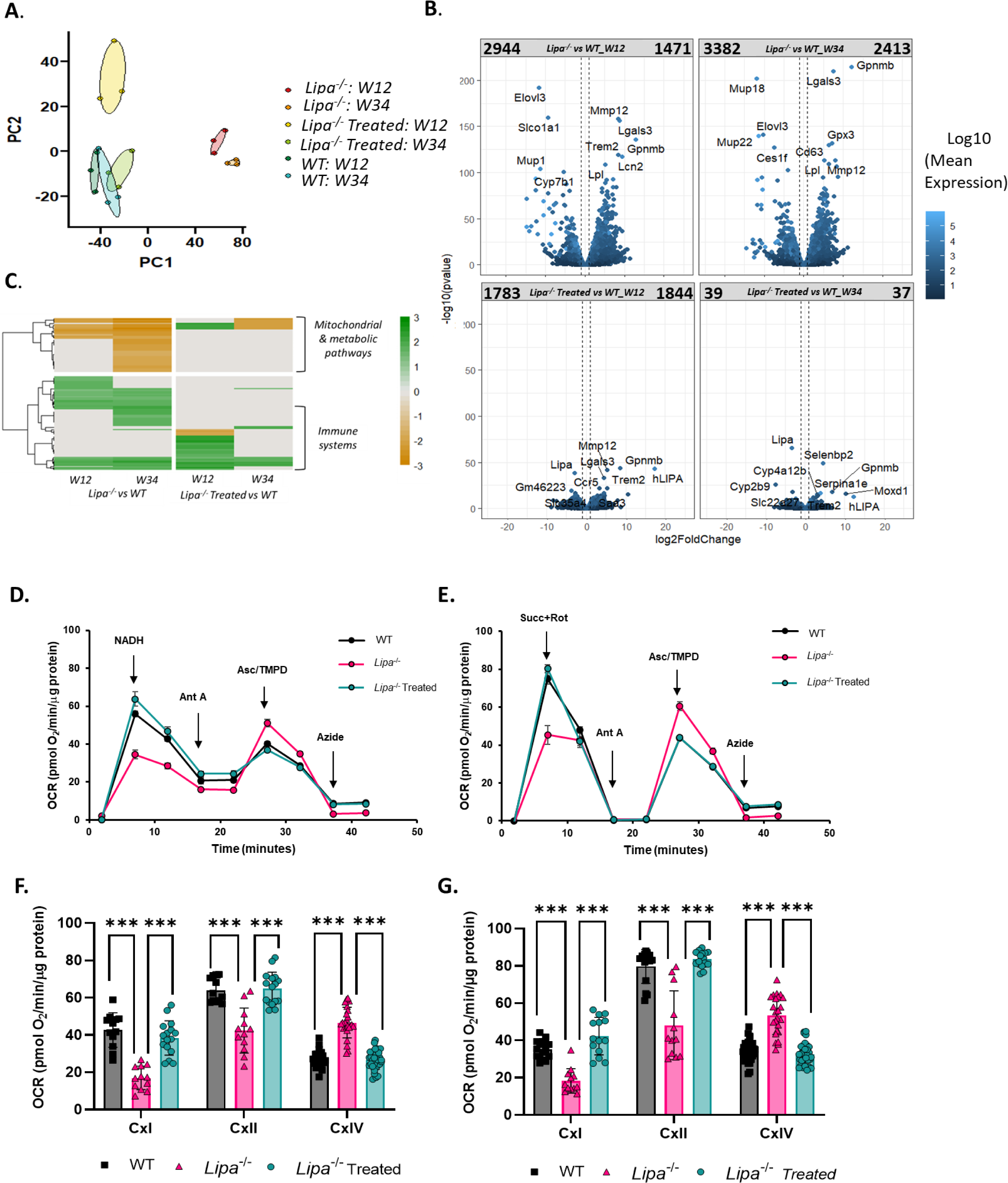
rAAV8 administration corrects expression of gene related to metabolic, immune, and mitochondria. Differential mRNA expression analysis was performed on liver WT, *Lipa*^-/-^ and rAAV8-treated *Lipa*^-/-^ mice. **(A)** Principal component analysis (PCA) visualizes the separation between experimental groups **(B)** Volcano plots display genes according to log2 fold change and -log10 P value. Colour shade indicates the log10 mean expression level of all analysed animals (n=3). For some dots the associated gene name is indicated. On top of each plot, the names indicate the samples compared, while the numbers indicate how many genes (dots) are present in each quadrant. Vertical dotted lines indicate ± 2-fold change. **(C)** Gene ontology analysis by Gene Set Enrichment Analysis (GSEA). Colour shades indicate normalized enrichment score (NES) of the indicated mice group comparison at 12- and 34-weeks post-injection. **(D** to **E)** Representative traces of the oxygen consumption rates (OCRs) in liver of WT, *Lipa*^-/-^ and rAAV8-treated *Lipa*^-/-^ mice at 12-weeks post-treatment. Where indicated, the following compounds were injected into the assay micro-chambers: NADH, Rotenone+Antimycin A (Rot+Ant A), Succinate+Rotenone (Succ+Rot), Antimycin A (AntA), Asc/TMPD (ascorbate+ TMPD), Azide. **(F** to **G.)** Metabolic parameters inferred from the OCR assays and corrected for residual activity in the presence of the respiratory chain inhibitors for mitochondria isolated from mouse at **(F)** 12- and **(G)** 34-weeks post injection. Bars indicate mean ± SD (n = 3-4 with 3 or 4 biological replicates for each group) and statistical significance was calculated using two-way ANOVA with Bonferroni post-hoc analysis (*p<0.033; **p<0.002; *** p<0.0002).

Differential gene expression analysis revealed that ∼4400 and ∼5800 genes changed between WT and *Lipa*^-/-^ mice at 12- and 34-weeks, respectively (Fig. 7.B top row). Upon rAAV8 mediated h*LIPA* expression (Fig. S7), both the number and the fold change of dysregulated genes was reduced, with only 76 genes differing between treated *Lipa*^-/-^ and WT mice at 34-weeks post-injection (Fig. 7.B bottom row).

Gene ontology analysis of differentially expressed genes between WT and *Lipa*^-/-^ mice identified two highly dysregulated pathways: (i) the downregulation of the metabolic and mitochondrial pathways in *Lipa*^-/-^ mice; (ii) the upregulation of the immune systems in *Lipa*^-/-^ mice (Fig. 7. C; left panel). rAAV8 administration normalized gene expression related to metabolic pathways at 12- and 34-weeks, while gene associated with the immune system were normalized only at 34-weeks (Fig. 7. C; right panel).

To confirm mitochondrial impairment, we isolated mitochondria from WT, *Lipa*^-/-^ and rAAV8-treated *Lipa*^-/-^ frozen mouse liver (at 12- and 34-weeks post-injection) and measured mitochondrial oxidative function. In particular, we assessed the efficiency of the electron transport chain complexes working together by exogenous substrates supplying electrons to complex I (NADH), II (succinate) and IV (ascorbate + N,N,N’,N’-tetramethyl-p-phenylenediamine dihydrochloride (TMPD)) (Fig 7. D-G). The oxygen consumption rates (OCR) elicited by complex I and II substrates (NADH and succinate respectively) were significantly reduced in *Lipa*^-/-^ mitochondria but completely restored upon rAAV8 administration already 12-weeks post-injection (Fig. 7.D-G). Surprisingly, the addition of complex IV substrates (ascorbate + TMPD) revealed a different scenario, with *Lipa*^-/-^ mitochondria showing a significant increase of respiratory activity compared to WT and rAAV8-treated *Lipa*^-/-^ mitochondria, probably due to a compensatory effect (*14*, *15*).

Altogether, these results suggest a reduced mitochondrial respiratory capacity of *Lipa*^-/-^ mitochondria, which was fully corrected by rAAV8 treatment.

Overall, administration of 3×10^12^ vg/kg of rAAV8 was sufficient to stably and completely correct the pathological phenotype of the *Lipa*^-/-^ mouse model for 34-weeks post-injection.

## DISCUSSION

In this study, we evaluated a liver directed gene therapy for LAL-D in a *Lipa*^-/-^ mouse model that displays reduced lifespan, failure to thrive and severe hepatosplenomegaly associated with lipid accumulation. By using a single-stranded rAAV8 encoding for the human *LIPA* cDNA under the control of the hepatocyte specific human α1-antitrypsin promoter, we demonstrated that transduced hepatocytes could express and secrete LAL, which retains its enzymatic activity and systemically cross-corrects LAL-D metabolic defect. In addition, we established the minimal rAAV8 vector dose required to extend mice survival, restore lipid homeostasis, and improve transcriptomic imbalance and mitochondrial activity.

We intravenously injected *Lipa*^-/-^ mice with four different rAAV8 doses and observed a transduction level and LAL activity proportional to the injected dose, suggesting that LAL overexpression does not result in liver toxicity. Secreted LAL was able to cross-correct blood cells and restore haematological parameters. In addition, expressed LAL ameliorated mouse weight and growth rate, corrected hepatosplenomegaly, and reduced lipid accumulation in the jejunum. Interestingly, even the lowest rAAV8 dose (5×10^11^ vg/kg) significantly reduced lipid accumulation and decreased disease burden.

We then performed a more detailed study using the minimal rAAV8 dose (3×10^12^ vg/kg) that improved lifespan and ameliorated both mouse weight and growth rate up to 34-week post-treatment. In particular, we evaluated short-term (4-weeks), middle term (12-weeks) and long-term (34-weeks) disease correction to confirm the efficacy and safety of our gene therapy treatment over time. Although we observed efficient hepatocyte transduction and LAL expression at all time points, there was a decrease between 4- and 12-weeks. Since rAAV genome remains mainly episomal, cellular proliferation during liver growth in young mice could explain this reduction (*16*). Interestingly, LAL expression fully corrected haematological parameters, dyslipidaemia and hepatic transaminase from 2-weeks post-injection and lipid storage and tissue architecture of liver, spleen, jejunum, and thymus from 4-weeks post-injection. Few small lipid droplets remained visible in the liver at late time points possibly due to transduction zonation (*17*) or reduced rAAV8 transduction in area of pre-existing liver injury in *Lipa*^-/-^ mice, e.g. fibrosis (*18*). Finally, by liver transcriptomic analysis and mitochondrial respirometry we observed a dysregulation of metabolic and immune system pathways (*19*) and an impairment in mitochondrial activity, which were fully normalized at 34-weeks post-injection. Overall, our *in vivo* gene therapy strategy appears to be an effective approach to achieve stable LAL expression to correct LAL-D phenotype in a relevant *Lipa^-/-^* mouse model. The current standard of treatment for patients affected by LAL-D is life-long weekly administration of recombinant human LAL enzyme (Sebelipase alfa, Kanuma) to reduce lipid storage and alleviate disease symptoms (*20*). Although effective in managing the condition, this treatment is not curative, and its effectiveness varies. Furthermore, it does not completely normalize all clinical manifestations, such as abdominal distortion, hepatosplenomegaly, and lipid accumulation in lymph nodes and the digestive wall (*21*). In addition, the repetitive administration of ERT may lead to the development of neutralising antibodies against the therapeutic enzyme, diminishing its efficacy over time (*22*). The described rAAV8 gene therapy has the potential to be a one-time curative treatment that maintains a constant level of therapeutic proteins and induces immune tolerance to the expressed transgene (*23*, *24*) or modulates an already existing immune response (*25–27*). Furthermore, hepatocyte-expressed LAL should result in enhanced pharmacokinetics and uptake of the enzyme in peripheral tissues compared to ERT, for which differential organ response is observed (*28*, *29*), and undergo endogenous post-translational modifications (e.g., glycosylation), which could reduce the development of neutralizing antibodies (*30*).

Lam *et al*. (*31*) recently described a liver directed gene therapy approach based on a recombinant self-complementary AAV vector serotype rh74 (rscAAVh74) encoding *LIPA* gene under the control of the ubiquitous mini cytomegalovirus promoter. Administration of the vector to *Lipa*^-/-^ mice successfully corrected the LAL-D phenotype by decreasing hepatosplenomegaly, dyslipidaemia, liver damage, and lipid accumulation. Although promising, this study has several limitations that requires further investigations. The authors used the rAAVh74 serotype, which has been selected to transduce muscles (*32–34*) and to treat muscular disorders (Dysferlinopathies: NCT02710500, Duchenne Muscular Dystrophy: NCT02704325, NCT02376816, NCT03375164). In addition, they used a self-complementary rAAV, which may increase the innate immune response compared to natural single-stranded rAAV vectors (*35–37*). Finally, to drive the expression of the LAL transgene they used the viral CMV promoter, which may become inactive due to DNA methylation (*38*, *39*). As an alternative, we propose using a single stranded rAAV8 to deliver the *LIPA* cDNA under the control of a liver-specific promoter. The rAAV8 serotype is currently the gold-standard for liver targeting and has been successfully tested in clinical trials for various metabolic disorders, such as glycogen storage disease type 1a (NCT03517085), Crigler Najjar (NCT03466463), mucopolysaccharidosis type VI (MPSVI) (NCT03173521), and Pompe disease (NCT03533673 and NCT04093349). It is worth noting that our rAAV8 corrective dose was ∼10 times lower than the previously suggested dose, reducing the risk of patient toxicity (*40*). Regarding the mouse model, the authors used a *Lipa*^-/-^ mouse model with normal body weight and growth rate and did not perform any survival analysis (*31*). Our *Lipa*^-/-^ mouse model (*11*), on a different genetic background and with a different LIPA mutation, allowed us to assess rAAV8-mediated correction of body weight, growth rate and lifespan. In addition, we conducted a follow-up analysis for up to 34-weeks post-injection and assessed biweekly several haematological parameters as a simple method to evaluate disease progression and gene therapy correction over time. Finally, using liver transcriptome analysis, we demonstrated a reduction in inflammation and correction of metabolic and mitochondrial pathways. To assess whether the mitochondrial alterations detected by transcriptomic analysis resulted in functional modifications, we measured the respiratory capacity of mitochondria. As reported in skeletal muscle fibers of *Lipa*^-/-^ mice (*41*) and in line with other lysosomal storage disorders (*42*), we found a significant decline of mitochondrial respiratory fluxes in mitochondria from *Lipa*^-/-^ livers. The respiratory fluxes were reduced when using Complex-I or Complex-II-dependent substrates, whereas they were increased when using exogenous substrates supplying electrons to Complex IV, suggesting a crucial role for Complex IV in compensating energy demand (*14*, *15*). Interestingly, administration of rAAV8 to *Lipa*^-/-^ mice fully restored the mitochondrial defect, both 12- and 34-weeks post-injection, confirming the effectiveness of our proposed therapy.

This *in vivo* gene therapy approach may still be hampered by preexisting neutralizing antibodies and cell-mediated immune responses against rAAV8 vectors with a 20-30% prevalence (*43*, *44*). A possible solution could be Imlifidase, a protease able to cleave IgG, that was shown to reduce pre-existing neutralizing antibody reactions against rAAV8 capsids in non-human primates, thus allowing vector re-administration (*45*). In addition, novel rAAV serotypes could be tested to reduce immunogenicity and increase liver targeting specificity (*46*, *47*), potentially reducing the required rAAV dose. To further decrease the rAAV dose and maximize therapeutic benefits, we will optimise codon usage and change the signal peptide sequence of the *LIPA* cDNA to improve LAL expression and secretion (*48*).

Although this rAAV8 strategy would be effective in patients with CESD, its efficacy would be limited in infants with WD, where episomal rAAV genome would be progressively diluted and lost upon liver growth (*49*, *16*). Therefore, a clinical translation will require treating patients with ERT until they reach the minimal age for AAV treatment (5 years) (*50*), similarly to what has recently been described for allogenic hematopoietic stem cell transplantation in LAL-D (*51*, *7*). In addition, research on haemophilia showed stable expression of therapeutic protein for up to 10-years using rAAV8 in human (*52*) and dog (*53*), possibly due to random genomic integration of rAAV DNA (*54*). Following this observation, we will investigate the use of DNA nucleases to achieve stable liver transgene targeted integration in a safe and highly transcribed genomic locus (*55–57*).

A possible alternative to rAAV would be a gene therapy strategy based on correction of autologous hematopoietic stem cells, which has already been successful for other lysosomal storage disorders (*58–61*). Extra caution will be required to avoid insertional mutagenesis, since LAL-D itself could be an initial trigger for tumour development (*62*).

Finally, to prepare for clinical trial, we plan to treat the larger rat *Lipa^-/-^* model that displays a severe disease phenotype closer to that of WD patients and has a life expectancy of 120 days (*63*).

In conclusion, our data suggest that a liver directed rAAV8 gene therapy strategy may be an effective curative treatment for LAL-D.

## MATERIALS AND METHODS

### Study design

This study was designed to evaluate the therapeutic potential of liver-based in vivo gene therapy using rAAV8.hAAT.LIPA for LAL-D. We selected an *Lipa^-/-^* mouse model that mimics the pathological lipids accumulation in the liver, spleen, and intestine, leading to reduced lifespan. To minimize the influence of sex on rAAV transduction (*64*) and immune reactions (*65*), male *Lipa^-/-^* mice were chosen for the study. We initially evaluated the minimal doses of rAAV8 able to correct hematological compartment, hepatosplenomegaly, lipid accumulation and tissues architecture. Each group consisted of four animals, which were monitored for 12-weeks post-administration. After identifying the most suitable doses of rAAV8, the corrective effects were evaluated at three different time points: 4-, 12, and 34-weeks post-administration. At each time point, a subset of mice was analyzed, including WT mice (3, 10 and 6 used respectively), *Lipa^-/-^* mice (4, 5 and 4 used respectively), and rAAV8.hAAT.LIPA-treated *Lipa^-/-^* mice (4, 6 and 6 used respectively). At the 34-weeks post-treatment, only one *Lipa^-/-^* mouse remained alive, while the others were sacrificed upon reaching predefined endpoints set by the ethical committee CEEA-51. Random assignment of mice to experimental groups was conducted to minimize bias.

### Mice

Heterozygous Lipa^tm1a(EUCOMM)Hmgu/Biat^ mice were crossed with heterozygous mice expressing the Cre recombinase under the control of the CMV promotor to delete exon 4 of the mouse *Lipa* gene (*11*) (Fig. S.1.A). Homozygous mice were then backcrossed on the C57/BL6N background for >8 generations. Mice were maintained in a specific pathogen-free (SPF) environment with a regular light-dark (12 h/12 h) cycle and *ad libitum* access to food (standard chow) and water. This study was approved by the ethical committee CEEA-51 and conducted according to French and European legislation on animal experimentation (APAFIS #33620-2021101416506495). Genotyping was performed by PCR on DNA extracted from the tail using 50 mM NaOH and Tris-HCl pH8. KAPA2G Fast ReadyMix (Kapa Biosystem, Wilmington, MA, USA) was used for PCR with using 0.65 µmol of each primer (Table 1, purchased at Thermo Fisher Scientific Waltham, MA) and 1µL of extracted DNA in a 12.5 µL reaction as per the manufacturer’s protocol. Genotyping bands were sequenced. Age and sex matched littermate controls were used for all experiments.

**Table 1).**
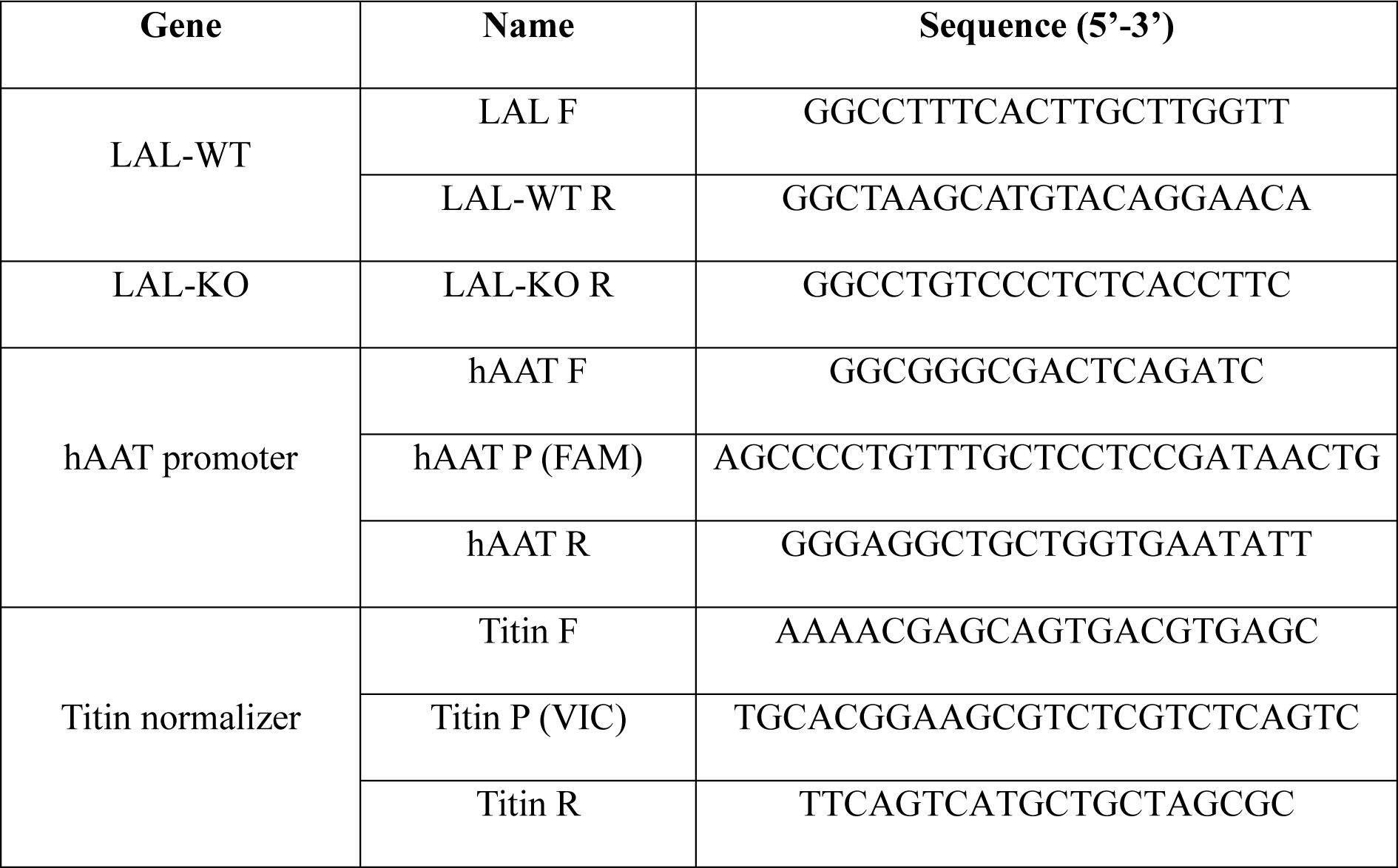
List of primers.

### Cloning and production of rAAV8

The cDNA of human *LIPA* gene (Gene ID:3988) with a C-terminal 3xHA tag (*55*) was cloned in a standard rAAV2/8 vector backbone under the control of the hAAT promoter and optimized HBB2 intron (*66*). Recombinant single-stranded rAAV2/8 was produced using an adenovirus-free triple transfection method of HEK293 and purified by single affinity chromatography (AVB Sepharose; GE Healthcare, Chicago, IL). The final product was formulated in sterile phosphate buffered saline containing 0.001% of Pluronic (F68; Sigma Aldrich, Saint Louis, MO), and stored at −80 °C (*67*, *68*).

### Administration of rAAV8

rAAV8.hAAT.LIPA vectors were administered via tail vein injection to 10- to 12-week-old *Lipa^-/-^* mice using an injection volume of 100µL. The animals’ weights were measured the day before administration, and viral solutions were prepared according to the animals’ weight. Mice were weighed weekly until sacrifice. Every two-weeks after injection, blood was drawn after isoflurane anaesthesia (IsoFlurin, Axience) from the retro-orbital veins using a heparin-coated capillary and collected into Eppendorf tube containing 3,8% citrate. At 4-, 12- and 34-weeks post injection, mice were sacrificed by cervical dislocation and their tissues (liver, spleen, jejunum, thymus) were divided in two parts. One part was snap-frozen in isopentane (−160C°) that was previously chilled in liquid nitrogen for histological analysis. The second part was disposed in 2mL bead tubes type D (Machery Nagel, Düren, Germany) and frozen in liquid nitrogen to assess rAAV8 biodistribution and to quantify lipid accumulation and LAL enzymatic activity.

### Blood and plasma analyses

The blood samples were analysed for standard haematological parameters (WBC counts and distribution, RBC count, haemoglobin, MCV and platelet counts, MPV) using an MS9.5 counter (Melet Schloessing, Cergy-Pontoise, France). After these analyses, blood was centrifuged for 10 min at 1,400 rcf and 4°C to recover plasma, which was stored at −80°C until further analysis. HDL, AST, and ALT levels were quantified in the plasma using specific colorimetric assays. A plasma control (Liquid assayed Multiqual; Bio-rad) was employed as a reference to quantify HDL, AST, and ALT. For the analysis, 10 μl of plasma were loaded onto specific chips designed for HDL, AST, and ALT measurements (HDL-C-P Ⅲ D and GPT/ALT-P Ⅲ; Fujifilm Europe Gmbh; FUJI DRI-CHEM NX500 - Sysmex). The concentration of HDL was measured in mg/dL, while ALT and AST were measured in U/L, providing quantitative data for these markers.

### Lipids analysis

Total lipids were extracted from 40 mg of frozen liver, spleen and jejunum using the chloroform-free Lipid Extraction Kit (Abcam, Waltham, MA, USA) according to the manufacturer’s protocol. Triglycerides were measured using the Triglycerides FS kit and total cholesterol was measured using the Cholesterol FS kit (both from DiaSys, Holzheim, Germany) following the manufacturer’s guidelines. Lipid concentrations were determined using a standard curve of triglycerides or cholesterol standards. Absorbance values were measured in duplicate using the EnSpire Multimode Plate Reader (Perkin Elmer, Waltham, MA, USA).

### Histological analysis

Serial 8 μm cross-sections of snap-frozen organs were cut on a Leica CM3050 S cryostat (Leica Biosystems). To minimise sampling errors, 2 sections of each specimen were obtained and stained with haematoxylin-eosin saffron (HES), Oil red O (ORO) and Sirius red (SR) according to standard procedures.

Images of ORO, HES and SR slides were digitised using an Axioscan Z1 slide scanner (Zeiss, Germany) with a Zeiss Plan-Apochromat 10X/0.45 M27 dry objective (ZEISS, Germany). Tile scan images were reconstructed with the ZEN software (ZEISS, Germany).

### ORO and SR quantification

ORO and SR images were processed using the pixel classification of QuPath 0.4.3 (*69*). For each staining and for each tissue, we created 2 classifiers, each time using several images for ground truth annotations. The first pixel classifier identifies pixels belonging to the tissue slice, excluding veins, folds, dust, bubbles, or other artifacts encountered, and draws an annotation of the analysable tissue. The second classifier was trained to identify fibrosis (for SR staining) or lipids (for ORO staining), based on manual ground truth annotations. This second classifier was then applied to the annotation created by the first classifier. The staining ratio for each tissue slice was obtained by dividing the total area of staining by the total surface area of the analysable tissue.

### LAL enzyme activity assay

Frozen liver, spleen and jejunum were homogenized in cold sterile PBS (Gibco) using FastPrep (MP Biomedical, Ohio, USA) followed by centrifugation for 5 min at 1,000xg and 4°C. The collected supernatant was frozen at −80°C prior to analysis. After sample processing, the frozen supernatant was centrifuged 20 min at 10,000×g to eliminate cell debris and the resulting supernatant was recovered. The protein content in the lysates was quantified using the BCA Protein Assay (Thermo Fisher Scientific). To determine LAL activity in liver and spleen, 1 µg of protein was used, whereas 10 µg of protein were used for the jejunum. Plasma was diluted 1/20 in sterile PBS prior to LAL activity measurement. Enzymatic activity was detected as previously described (*55*). Briefly, samples were incubated for 10 min at 37 °C with 42 μM Lalistat-2 (Sigma-Aldrich, St. Louis, MI, USA) or water and then transferred to an Optiplate 96 F plate (PerkinElmer). Fluorometric reactions were initiated with 75 μL of substrate buffer (340 μM 4-MUP, 0.9% Triton X-100 and 220 μM cardiolipin in 135mM acetate buffer pH 4.0). After 10 min, fluorescence was recorded (35 cycles, 30 sec intervals, 37 °C) using the SPARK TECAN Reader (Tecan, Austria). Kinetic parameters (average rate) were calculated using Magellan Software. LAL activity was quantified using this formula:

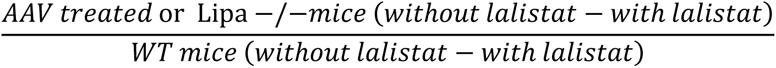

### Western Blot

Protein was extracted from frozen liver in sterile PBS using Fastprep crushing (MP Biomedical, Ohio, USA), followed by centrifugation for 10 min at 14,000×g and4°C to collect the supernatant. The protein content was quantified using BCA Protein Assay (Thermo Fisher Scientific) and 40 ng of protein were loaded as previously described (*55*). α-actinin was used as loading control and antibodies are referred in Table 2. The blots were imaged at 169 μm with the Odyssey imager and analysed with ImageStudio Lite software (Li-Cor Biosciences, Lincoln, NE, USA). After image background subtraction (average method, top/bottom), the band intensities were quantified and normalised with α-actinin signal.

**Table 2).**
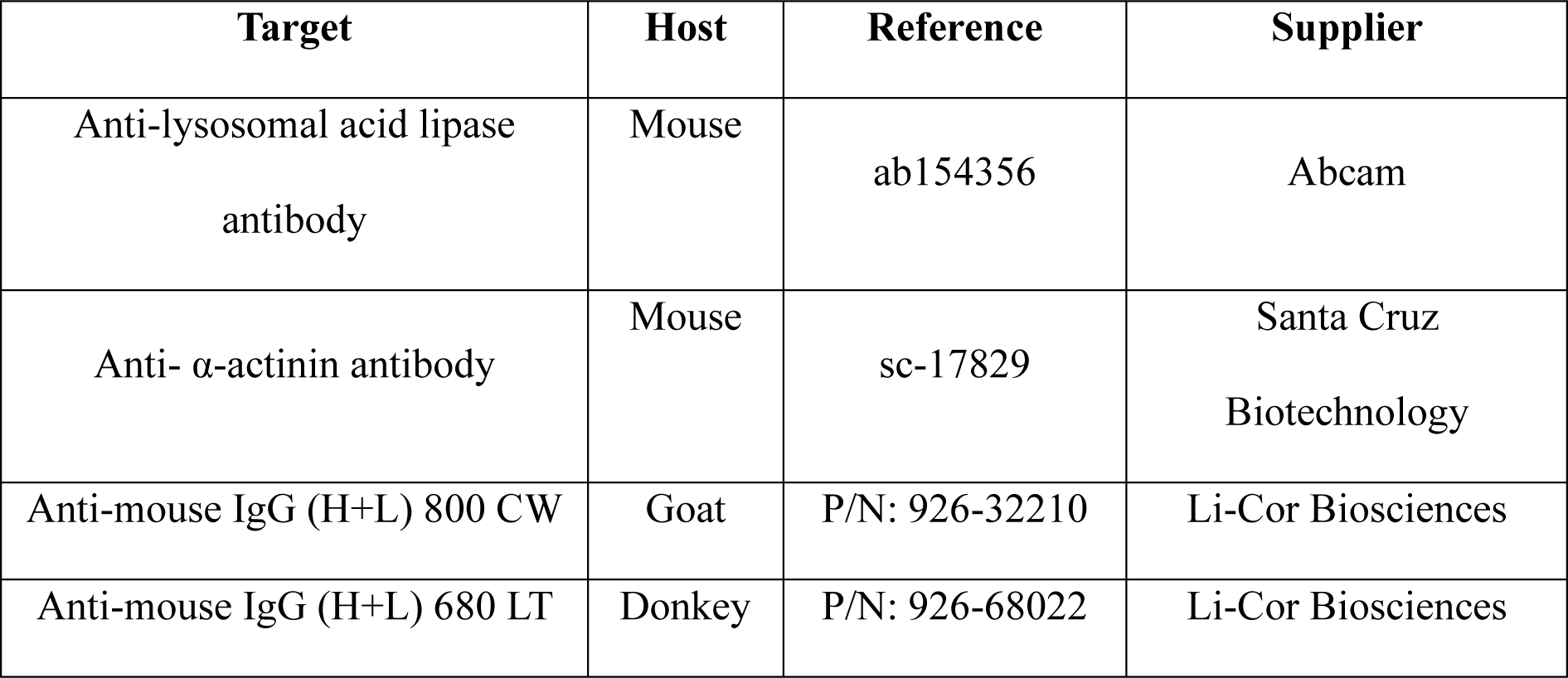
List of antibodies.

### Vector genomes

The genomic DNA was extracted from frozen organs using the DNA extraction kit (NucleoMag Pathogen; Macherey Nagel) and the KingFisher™ Flex Purification System with 96 Deep-Well Head (Thermo Fisher Scientific) following the manufacturer’s instructions. The number of vector copies per diploid genome was determined using Taqman qPCR with specific primers to amplify the hAAT promoter sequence with 100ng of extracted DNA. Titin was used as normalizer to determine the number of genome copies. Data were expressed as vector genome copies per diploid genome.

### RNA seq

Azenta Life Science (Leipzig, Germany) received 40 mg of frozen liver for RNA sequencing. The paired-end reads (2×150bp) were aligned to the reference transcriptome (GRCm39/ensemble cDNA release 110) using SALMON v1.9.0 (*70*). Default parameters were used in addition to arguments -l A and –validate Mappings. The quantification files were processed using R (4.3.0) and the tximport package (1.28.0) to perform pairwise group differential expression analysis using DESeq2 (1.40.1) on genes with > 100 reads across all samples (lfcThreshold = 0, pAdjustMethod = “fdr”, independentFiltering = T) (*71*). Pathway enrichment analysis and GSEA were performed using the reactomePA (1.44.0) and clusterProfiler (4.8.1) (*72*) packages for reactome (*73*) and GO terms respectively. Genes were considered dysregulated if the absolute Log fold change was >1 and the FDR-adjusted P value < 0.05. The RNA-seq data, both raw and processed, are available on the GEO data set: GSE252742.

### Metabolic fluxes analysis in mouse liver

Mitochondria were isolated from frozen liver of mouse at 12- and 34-weeks post injection, as previously described (*74*). Briefly, frozen liver tissue was minced and homogenized in MAS buffer containing 70 mM sucrose, 220 mM mannitol, 5 mM KH_2_PO_4_, 5 mM MgCl_2_, 1 mM EGTA and 2 mM HEPES pH 7.4. The homogenate was centrifuged at 1,000 × g for 5 min. The resulting supernatant was centrifuged at 10,000 × g for 10 min and the pellet was then resuspended in the same buffer. All centrifugation steps were carried out at 0–4 °C. Mitochondrial protein concentrations were determined using the Bradford method with bovine serum albumin as standard. The respiratory activity of freshly prepared mitochondria was measured with the XF96 extracellular flux analyser (Seahorse Bioscence, Agilent). Mitochondria (100 µg) were loaded into a Seahorse XF96 microplate in 20 μl of MAS. The loaded plate was centrifuged at 2,000 g for 5 min at 4°C (no brake) and an additional 130 μl of MAS containing cytochrome c (10 μg/ml, final concentration). The substrate was added as follows: NADH (2 mM), or succinate + rotenone (5 mM + 2 μM) was injected at port A; rotenone (2 µM) or rotenone + antimycin A (2 μM + 4 μM) at port B; TMPD + ascorbic acid (0.5 mM + 1 mM) at port C; and azide (50 mM) at port D. These conditions allowed for the determination of the respiratory capacity of mitochondria through Complex I, Complex II, and Complex IV.

### Statistical analysis

Statistical analyses were performed using GraphPad Prism version 9.00 for Windows (GraphPad Software, La Jolla, CA, USA, “www.graphpad.com”). One-way or two-way analysis of variance (ANOVA) with Tukey’s test were performed as indicated (alpha = 0.05). Values are expressed as mean ± standard deviation (SD). All unmarked comparisons reflect non-significant (p > 0.05) differences.

## List of Supplementary Materials

Fig. S1. Description of the *Lipa*^-/-^ mouse model.

Fig. S2. Blood analysis of *Lipa*^-/-^ mice treated with 4 different rAAV8 doses.

Fig. S3. Quantification of cholesterol and triglycerides per mg of liver, spleen, and jejunum 12-weeks post-injection.

Fig. S4. Long-term blood analysis of rAAV8.hAAT.LIPA treated *Lipa*^-/-^ mice.

Fig. S5. Quantification of cholesterol and triglycerides per mg of liver, spleen, and jejunum. Fig. S6. Quantification of lipid deposits in thymus.

Fig. S7. Read count of murine (m) and human (h) LIPA mRNA at 34-week post-injection.

## Acknowledgement

We thank Genethon “bioexperimentation team” for technical assistance on mice experimentation, and Genethon “histology platform” for providing histological section. The authors are Genopole’s members, first french biocluster dedicated to genetic, biotechnologies and biotherapies. We are grateful to the “Imaging and Cytometry Core Facility” of Genethon for technical support, member of the the National Infrastructure in Biology and Health France-BioImaging, to Ile-de-France Region, to Conseil Départemental de l’Essonne (ASTRE), INSERM and GIP Genopole, Evry for the purchase of the equipments. We thank Genethon “Vector Core Facility” and Dr. Bérangère Bertin for rAAV production and Pauline Vidal for technical support. We gratefully acknowledge the Conseil Général de l’Essonne (ASTRES) and Genopole Research in Evry for the financial help to purchase equipment. We thank the whole Amendola’s laboratory and Dr. François-Xavier Mauvais for fruitful discussion.

## Author contribution

M.L. conceived the study, designed, and performed experiments, analysed data, and wrote the manuscript. R.H., C.J. and J.O. performed experiments and analysed data. S.J. and J.C. performed histological staining acquisition and quantification. G.C. performed bioinformatic analysis of transcriptomic data. C.P. and C.S. conceived and F.L. and A.F. performed mitochondrial studies. N.V. and A.B. provided expertise. L.V.W. performed mice experimentation. G.R. provided the rAAV8 backbone plasmid and expertise. D.K. provided the *Lipa*^-/-^ mouse model and expertise. M.A. conceived the study, designed experiments, acquired funding, analysed data, and wrote the manuscript.

## Competitive interest

All authors declare no competing interests.

## References

1. W. V. Brown, R. J. Desnick, G. A. Grabowski, JCL Roundtable: enzyme replacement therapy for lipid storage disorders. J. Clin. Lipidol. 8, 463–472 (2014).

2. A. A. Aqul, C. M. Ramirez, A. M. Lopez, D. K. Burns, J. J. Repa, S. D. Turley, Molecular markers of brain cholesterol homeostasis are unchanged despite a smaller brain mass in a mouse model of cholesteryl ester storage disease. Lipids 57, 3–16 (2022).

3. M. Balwani, W. Balistreri, L. D’Antiga, J. Evans, E. Ros, F. Abel, D. P. Wilson, Lysosomal acid lipase deficiency manifestations in children and adults: Baseline data from an international registry. Liver Int. **n/a** (2023).

4. C. da R. Witeck, A. C. Schmitz, J. M. D. de Oliveira, A. L. Porporatti, G. D. L. Canto, M. M. de S. Pires, Lysosomal acid lipase deficiency in pediatric patients: a scoping review. J. Pediatr. (Rio J.) 98, 04–14 (2022).

5. J. Tolar, A. Petryk, K. Khan, K. Bjoraker, J. Jessurun, M. Dolan, T. Kivisto, L. Charnas, E. Shapiro, P. Orchard, Long-term metabolic, endocrine, and neuropsychological outcome of hematopoietic cell transplantation for Wolman disease. Bone Marrow Transplant. 43, 21 (2009).

6. D. L. Bernstein, S. Lobritto, A. Iuga, H. Remotti, T. Schiano, M. I. Fiel, M. Balwani, Lysosomal acid lipase deficiency allograft recurrence and liver failure-clinical outcomes of 18 liver transplantation patients. Mol. Genet. Metab. 124, 11–19 (2018).

7. J. E. Potter, G. Petts, A. Ghosh, F. J. White, J. L. Kinsella, S. Hughes, J. Roberts, A. Hodgkinson, K. Brammeier, H. Church, C. Merrigan, J. Hughes, P. Evans, H. Campbell, D. Bonney, W. G. Newman, B. W. Bigger, A. Broomfield, S. A. Jones, R. F. Wynn, Enzyme replacement therapy and hematopoietic stem cell transplant: a new paradigm of treatment in Wolman disease. Orphanet J. Rare Dis. 16, 235 (2021).

8. B. K. Burton, F. Feillet, K. N. Furuya, S. Marulkar, M. Balwani, Sebelipase alfa in children and adults with lysosomal acid lipase deficiency: Final results of the ARISE study. J. Hepatol. 76, 577– 587 (2022).

9. S. Vijay, A. Brassier, A. Ghosh, S. Fecarotta, F. Abel, S. Marulkar, S. A. Jones, “Long-Term Survival With Sebelipase Alfa Enzyme Replacement Therapy in Infants With Rapidly Progressive Lysosomal Acid Lipase Deficiency Final Results From 2 Open-Label Studies” (preprint, In Review, 2020); 10.21203/rs.3.rs-45422/v2.

10. H. Du, M. Heur, D. P. Witte, D. Ameis, G. A. Grabowski, Lysosomal Acid Lipase Deficiency: Correction of Lipid Storage by Adenovirus-Mediated Gene Transfer in Mice. Hum. Gene Ther. 13, 1361–1372 (2002).

11. C. Leopold, M. Duta-Mare, V. Sachdev, M. Goeritzer, L. K. Maresch, D. Kolb, H. Reicher, B. Wagner, T. Stojakovic, T. Ruelicke, G. Haemmerle, G. Hoefler, W. Sattler, D. Kratky, Hepatocyte-specific lysosomal acid lipase deficiency protects mice from diet-induced obesity but promotes hepatic inflammation. Biochim. Biophys. Acta BBA - Mol. Cell Biol. Lipids 1864, 500–511 (2019).

12. P. Qu, W. C. Shelley, M. C. Yoder, L. Wu, H. Du, C. Yan, Critical Roles of Lysosomal Acid Lipase in Myelopoiesis. Am. J. Pathol. 176, 2394–2404 (2010).

13. S. Schlager, N. Vujic, M. Korbelius, M. Duta-Mare, J. Dorow, C. Leopold, S. Rainer, M. Wegscheider, H. Reicher, U. Ceglarek, W. Sattler, B. Radovic, D. Kratky, Lysosomal lipid hydrolysis provides substrates for lipid mediator synthesis in murine macrophages. Oncotarget 8, 40037– 40051 (2017).

14. S. Licht-Mayer, G. R. Campbell, A. R. Mehta, K. McGill, A. Symonds, S. Al-Azki, G. Pryce, S. Zandee, C. Zhao, M. Kipp, K. J. Smith, D. Baker, D. Altmann, S. M. Anderton, Y. S. Kap, J. D. Laman, B. A. ’t Hart, M. Rodriguez, R. J. M. Franklin, S. Chandran, H. Lassmann, B. D. Trapp, D. J. Mahad, Axonal response of mitochondria to demyelination and complex IV activity within demyelinated axons in experimental models of multiple sclerosis. Neuropathol. Appl. Neurobiol. 49, e12851 (2023).

15. M. K. Tripathy, D. Mitra, Differential modulation of mitochondrial OXPHOS system during HIV-1 induced T-cell apoptosis: up regulation of Complex-IV subunit COX-II and its possible implications. Apoptosis Int. J. Program. Cell Death 15, 28–40 (2010).

16. L. Wang, H. Wang, P. Bell, D. McMenamin, J. M. Wilson, Hepatic gene transfer in neonatal mice by adeno-associated virus serotype 8 vector. Hum. Gene Ther. 23, 533–539 (2012).

17. P. Bell, L. Wang, G. Gao, M. E. Haskins, A. F. Tarantal, R. J. McCarter, Y. Zhu, H. Yu, J. M. Wilson, Inverse zonation of hepatocyte transduction with AAV vectors between mice and non-human primates. Mol. Genet. Metab. 104, 395–403 (2011).

18. A. Cantore, C. Simoni, J. Nozi, F. Starinieri, T. L. Bella, E. Manta, C. Negri, M. Biffi, R. Norata, M. Rocchi, F. Sanvito, G. Ronzitti, E. Barbon, Liver fibrosis negatively impacts in vivo gene transfer to hepatocytes. [Preprint] (2024). 10.21203/rs.3.rs-4011604/v1.

19. I. Bradić, L. Liesinger, K. B. Kuentzel, N. Vujić, M. Trauner, R. Birner-Gruenberger, D. Kratky, Metabolic changes and propensity for inflammation, fibrosis, and cancer in livers of mice lacking lysosomal acid lipase. J. Lipid Res., 100427 (2023).

20. T. Demaret, F. Lacaille, C. Wicker, J.-B. Arnoux, J. Bouchereau, C. Belloche, C. Gitiaux, D. Grevent, C. Broissand, D. Adjaoud, M.-T. Abi Warde, D. Plantaz, S. Bekri, P. de Lonlay, A. Brassier, Sebelipase alfa enzyme replacement therapy in Wolman disease: a nationwide cohort with up to ten years of follow-up. Orphanet J. Rare Dis. 16, 1–9 (2021).

21. W. B. Hannah, K. Ryan, S. Pendyal, T. A. Burrow, S. E. Harley, M. Cordell, C. M. McCall, A. M. Mavis, Q. K.-G. Tan, P. S. Kishnani, Clinical insights from Wolman disease: Evaluating infantile hepatosplenomegaly. Am. J. Med. Genet. A. 188, 3364–3368 (2022).

22. J. Wang, J. Lozier, G. Johnson, S. Kirshner, D. Verthelyi, A. Pariser, E. Shores, A. Rosenberg, Neutralizing antibodies to therapeutic enzymes: considerations for testing, prevention and treatment. Nat. Biotechnol. 26, 901–908 (2008).

23. G. D. Keeler, D. M. Markusic, B. E. Hoffman, Liver induced transgene tolerance with AAV vectors. Cell. Immunol. 342, 103728 (2019).

24. P. A. LoDuca, B. E. Hoffman, R. W. Herzog, Hepatic Gene Transfer as a Means of Tolerance Induction to Transgene Products. Curr. Gene Ther. 9, 104–114 (2009).

25. S. Han, G. Ronzitti, B. Arnson, C. Leborgne, S. Li, F. Mingozzi, D. Koeberl, Low-dose liver-targeted gene therapy for Pompe disease enhances therapeutic efficacy of ERT via immune tolerance induction. Mol. Ther.-Methods Clin. Dev. 4, 126–136 (2017).

26. D. M. Markusic, B. E. Hoffman, G. Q. Perrin, S. Nayak, X. Wang, P. A. LoDuca, K. A. High, R. W. Herzog, Effective gene therapy for haemophilic mice with pathogenic factor IX antibodies. EMBO Mol. Med. 5, 1698–1709 (2013).

27. B. Sun, A. Bird, S. P. Young, P. S. Kishnani, Y.-T. Chen, D. D. Koeberl, Enhanced response to enzyme replacement therapy in Pompe disease after the induction of immune tolerance. Am. J. Hum. Genet. 81, 1042–1049 (2007).

28. Y. Sun, Y.-H. Xu, H. Du, B. Quinn, B. Liou, L. Stanton, V. Inskeep, H. Ran, P. Jakubowitz, N. Grilliot, G. A. Grabowski, Reversal of advanced disease in lysosomal acid lipase deficient mice: A model for lysosomal acid lipase deficiency disease. Mol. Genet. Metab. 112, 229–241 (2014).

29. H. Costa-Verdera, F. Collaud, C. R. Riling, P. Sellier, J. M. L. Nordin, G. M. Preston, U. Cagin, J. Fabregue, S. Barral, M. Moya-Nilges, J. Krijnse-Locker, L. van Wittenberghe, N. Daniele, B. Gjata, J. Cosette, C. Abad, M. Simon-Sola, S. Charles, M. Li, M. Crosariol, T. Antrilli, W. J. Quinn, D. A. Gross, O. Boyer, X. M. Anguela, S. M. Armour, P. Colella, G. Ronzitti, F. Mingozzi, Hepatic expression of GAA results in enhanced enzyme bioavailability in mice and non-human primates. Nat. Commun. 12, 6393 (2021).

30. R. Jefferis, Posttranslational Modifications and the Immunogenicity of Biotherapeutics. J. Immunol. Res. 2016, 5358272 (2016).

31. P. Lam, A. Ashbrook, D. A. Zygmunt, C. Yan, H. Du, P. T. Martin, Therapeutic efficacy of rscAAVrh74.miniCMV.LIPA gene therapy in a mouse model of lysosomal acid lipase deficiency. Mol. Ther. - Methods Clin. Dev. 26, 413–426 (2022).

32. E. R. Pozsgai, D. A. Griffin, K. N. Heller, J. R. Mendell, L. R. Rodino-Klapac, Systemic AAV-Mediated β-Sarcoglycan Delivery Targeting Cardiac and Skeletal Muscle Ameliorates Histological and Functional Deficits in LGMD2E Mice. Mol. Ther. 25, 855–869 (2017).

33. J. Shoti, K. Qing, G. D. Keeler, D. Duan, B. J. Byrne, A. Srivastava, Development of capsid- and genome-modified optimized AAVrh74 vectors for muscle gene therapy. Mol. Ther. - Methods Clin. Dev., 101147 (2023).

34. P. C. Sondergaard, D. A. Griffin, E. R. Pozsgai, R. W. Johnson, W. E. Grose, K. N. Heller, K. M. Shontz, C. L. Montgomery, J. Liu, K. R. Clark, Z. Sahenk, J. R. Mendell, L. R. Rodino-Klapac, AAV.Dysferlin Overlap Vectors Restore Function in Dysferlinopathy Animal Models. Ann. Clin. Transl. Neurol. 2, 256–270 (2015).

35. A. T. Martino, M. Suzuki, D. M. Markusic, I. Zolotukhin, R. C. Ryals, B. Moghimi, H. C. J. Ertl, D. A. Muruve, B. Lee, R. W. Herzog, The genome of self-complementary adeno-associated viral vectors increases Toll-like receptor 9-dependent innate immune responses in the liver. Blood 117, 6459–6468 (2011).

36. T. Wu, K. Töpfer, S.-W. Lin, H. Li, A. Bian, X. Y. Zhou, K. A. High, H. C. Ertl, Self-complementary AAVs Induce More Potent Transgene Product-specific Immune Responses Compared to a Single-stranded Genome. Mol. Ther. 20, 572–579 (2012).

37. J. Zhu, X. Huang, Y. Yang, The TLR9-MyD88 pathway is critical for adaptive immune responses to adeno-associated virus gene therapy vectors in mice. J. Clin. Invest. 119, 2388–2398 (2009).

38. A. R. Brooks, R. N. Harkins, P. Wang, H. S. Qian, P. Liu, G. M. Rubanyi, Transcriptional silencing is associated with extensive methylation of the CMV promoter following adenoviral gene delivery to muscle. J. Gene Med. 6, 395–404 (2004).

39. S. Prösch, J. Stein, K. Staak, C. Liebenthal, H. D. Volk, D. H. Krüger, Inactivation of the very strong HCMV immediate early promoter by DNA CpG methylation in vitro. Biol. Chem. Hoppe. Seyler 377, 195–201 (1996).

40. High-dose AAV gene therapy deaths. Nat. Biotechnol. 38, 910 (2020).

41. A. Akhmetshina, V. Bianco, I. Bradić, M. Korbelius, A. Pirchheim, K. B. Kuentzel, T. O. Eichmann, H. Hinteregger, D. Kolb, H. Habisch, L. Liesinger, T. Madl, W. Sattler, B. Radović, S. Sedej, R. Birner-Gruenberger, N. Vujić, D. Kratky, Loss of lysosomal acid lipase results in mitochondrial dysfunction and fiber switch in skeletal muscles of mice. Mol. Metab. 79, 101869 (2024).

42. K. M. Stepien, F. Roncaroli, N. Turton, C. J. Hendriksz, M. Roberts, R. A. Heaton, I. Hargreaves, Mechanisms of Mitochondrial Dysfunction in Lysosomal Storage Disorders: A Review. J. Clin. Med. 9, 2596 (2020).

43. A. Kruzik, D. Fetahagic, B. Hartlieb, S. Dorn, H. Koppensteiner, F. M. Horling, F. Scheiflinger, B. M. Reipert, M. de la Rosa, Prevalence of Anti-Adeno-Associated Virus Immune Responses in International Cohorts of Healthy Donors. Mol. Ther. Methods Clin. Dev. 14, 126–133 (2019).

44. R. Xicluna, A. Avenel, C. Vandamme, M. Devaux, N. Jaulin, C. Couzinie, J. L. Duff, A. Charrier, M. Guilbaud, O. Adjali, G. Gernoux, Prevalence study of cellular capsid-specific immune responses to AAV1, 2, 4, 5, 8, 9 and rh10 reveals particular features for AAV9. bioRxiv [Preprint] (2023). 10.1101/2023.12.20.570598.

45. C. Leborgne, E. Barbon, J. M. Alexander, H. Hanby, S. Delignat, D. M. Cohen, F. Collaud, S. Muraleetharan, D. Lupo, J. Silverberg, K. Huang, L. van Wittengerghe, B. Marolleau, A. Miranda, A. Fabiano, V. Daventure, H. Beck, X. M. Anguela, G. Ronzitti, S. M. Armour, S. Lacroix-Desmazes, F. Mingozzi, IgG-cleaving endopeptidase enables in vivo gene therapy in the presence of anti-AAV neutralizing antibodies. Nat. Med. 26, 1096–1101 (2020).

46. L. Lisowski, A. P. Dane, K. Chu, Y. Zhang, S. C. Cunningham, E. M. Wilson, S. Nygaard, M. Grompe, I. E. Alexander, M. A. Kay, Selection and evaluation of clinically relevant AAV variants in a xenograft liver model. Nature 506, 382–386 (2014).

47. K. Vercauteren, B. E. Hoffman, I. Zolotukhin, G. D. Keeler, J. W. Xiao, E. Basner-Tschakarjan, K. A. High, H. C. Ertl, C. M. Rice, A. Srivastava, Y. P. de Jong, R. W. Herzog, Superior In vivo Transduction of Human Hepatocytes Using Engineered AAV3 Capsid. Mol. Ther. 24, 1042–1049 (2016).

48. F. Puzzo, P. Colella, M. G. Biferi, D. Bali, N. K. Paulk, P. Vidal, F. Collaud, M. Simon-Sola, S. Charles, R. Hardet, C. Leborgne, A. Meliani, M. Cohen-Tannoudji, S. Astord, B. Gjata, P. Sellier, L. van Wittenberghe, A. Vignaud, F. Boisgerault, M. Barkats, P. Laforet, M. A. Kay, D. D. Koeberl, G. Ronzitti, F. Mingozzi, Rescue of Pompe disease in mice by AAV-mediated liver delivery of secretable acid α-glucosidase. Sci. Transl. Med. 9, eaam6375 (2017).

49. S. C. Cunningham, A. P. Dane, A. Spinoulas, I. E. Alexander, Gene Delivery to the Juvenile Mouse Liver Using AAV2/8 Vectors. Mol. Ther. J. Am. Soc. Gene Ther. 16, 1081–1088 (2008).

50. N. Brunetti-Pierri, R. Ferla, V. M. Ginocchio, A. Rossi, S. Fecarotta, R. Romano, G. Parenti, Y. Yildiz, S. Zancan, V. Pecorella, M. Dell’Anno, M. Graziano, M. Alliegro, G. Andria, F. Santamaria, R. Brunetti-Pierri, F. Simonelli, V. Nigro, M. Vargas, G. Servillo, F. Borgia, E. Soscia, M. Gargaro, S. Funghini, N. Tedesco, P. R. Le Brun, C. A. Rupar, C. Prasad, M. O’Callaghan, J. J. Mitchell, O. Danos, J.-B. Marteau, S. Galimberti, M. G. Valsecchi, P. Veron, F. Mingozzi, F. Fallarino, G. la Marca, H. S. Sivri, A. Auricchio, Liver-Directed Adeno-Associated Virus-Mediated Gene Therapy for Mucopolysaccharidosis Type VI. NEJM Evid. 1, EVIDoa2200052 (2022).

51. S. K. Eskandari, E. G. M. Revenich, D. J. Pot, F. de Boer, M. Bierings, F. J. van Spronsen, P. M. van Hasselt, C. A. Lindemans, C. M. A. Lubout, High-Dose ERT, Rituximab, and Early HSCT in an Infant with Wolman’s Disease. N. Engl. J. Med. 390, 623–629 (2024).

52. A. C. Nathwani, U. Reiss, E. Tuddenham, P. Chowdary, J. McIntosh, A. Riddell, J. Pie, J. N. Mahlangu, M. Recht, Y.-M. Shen, K. G. Halka, M. M. Meagher, A. W. Nienhuis, A. M. Davidoff, S. Mangles, C. L. Morton, Z. Junfang, V. C. Radulescu, Adeno-Associated Mediated Gene Transfer for Hemophilia B:8 Year Follow up and Impact of Removing “Empty Viral Particles” on Safety and Efficacy of Gene Transfer. Blood 132, 491 (2018).

53. P. Batty, A. M. Mo, D. Hurlbut, J. Ishida, B. Yates, C. Brown, L. Harpell, C. Hough, A. Pender, E. K. Rimmer, S. Sardo Infirri, A. Winterborn, S. Fong, D. Lillicrap, Long-term follow-up of liver-directed, adeno-associated vector-mediated gene therapy in the canine model of hemophilia A. Blood 140, 2672–2683 (2022).

54. J. A. Greig, K. M. Martins, C. Breton, R. J. Lamontagne, Y. Zhu, Z. He, J. White, J.-X. Zhu, J. A. Chichester, Q. Zheng, Z. Zhang, P. Bell, L. Wang, J. M. Wilson, Integrated vector genomes may contribute to long-term expression in primate liver after AAV administration. Nat. Biotechnol., 1–11 (2023).

55. G. Pavani, M. Laurent, A. Fabiano, E. Cantelli, A. Sakkal, G. Corre, P. J. Lenting, J.-P. Concordet, M. Toueille, A. Miccio, M. Amendola, Ex vivo editing of human hematopoietic stem cells for erythroid expression of therapeutic proteins. Nat. Commun. 11, 3778 (2020).

56. G. Pavani, M. Amendola, Targeted Gene Delivery: Where to Land. Front. Genome Ed. 2, 609650 (2020).

57. R. Sharma, X. M. Anguela, Y. Doyon, T. Wechsler, R. C. DeKelver, S. Sproul, D. E. Paschon, J. C. Miller, R. J. Davidson, D. Shivak, S. Zhou, J. Rieders, P. D. Gregory, M. C. Holmes, E. J. Rebar, K. A. High, In vivo genome editing of the albumin locus as a platform for protein replacement therapy. Blood 126, 1777–1784 (2015).

58. F. Fumagalli, V. Calbi, M. G. Natali Sora, M. Sessa, C. Baldoli, P. M. V. Rancoita, F. Ciotti, M. Sarzana, M. Fraschini, A. A. Zambon, S. Acquati, D. Redaelli, V. Attanasio, S. Miglietta, F. De Mattia, F. Barzaghi, F. Ferrua, M. Migliavacca, F. Tucci, V. Gallo, U. Del Carro, S. Canale, I. Spiga, L. Lorioli, S. Recupero, E. S. Fratini, F. Morena, P. Silvani, M. R. Calvi, M. Facchini, S. Locatelli, A. Corti, S. Zancan, G. Antonioli, G. Farinelli, M. Gabaldo, J. Garcia-Segovia, L. C. Schwab, G. F. Downey, M. Filippi, M. P. Cicalese, S. Martino, C. Di Serio, F. Ciceri, M. E. Bernardo, L. Naldini, A. Biffi, A. Aiuti, Lentiviral haematopoietic stem-cell gene therapy for early-onset metachromatic leukodystrophy: long-term results from a non-randomised, open-label, phase 1/2 trial and expanded access. The Lancet 399, 372–383 (2022).

59. B. Gentner, F. Tucci, S. Galimberti, F. Fumagalli, M. De Pellegrin, P. Silvani, C. Camesasca, S. Pontesilli, S. Darin, F. Ciotti, M. Sarzana, G. Consiglieri, C. Filisetti, G. Forni, L. Passerini, D. Tomasoni, D. Cesana, A. Calabria, G. Spinozzi, M.-P. Cicalese, V. Calbi, M. Migliavacca, F. Barzaghi, F. Ferrua, V. Gallo, S. Miglietta, E. Zonari, P. S. Cheruku, C. Forni, M. Facchini, A. Corti, M. Gabaldo, S. Zancan, S. Gasperini, A. Rovelli, J.-J. Boelens, S. A. Jones, R. Wynn, C. Baldoli, E. Montini, S. Gregori, F. Ciceri, M. G. Valsecchi, G. la Marca, R. Parini, L. Naldini, A. Aiuti, M.-E. Bernardo, Hematopoietic Stem-and Progenitor-Cell Gene Therapy for Hurler Syndrome. N. Engl. J. Med. 385, 1929–1940 (2021).

60. F. Eichler, C. Duncan, P. L. Musolino, P. J. Orchard, S. De Oliveira, A. J. Thrasher, M. Armant, C. Dansereau, T. C. Lund, W. P. Miller, G. V. Raymond, R. Sankar, A. J. Shah, C. Sevin, H. B. Gaspar, P. Gissen, H. Amartino, D. Bratkovic, N. J. C. Smith, A. M. Paker, E. Shamir, T. O’Meara, D. Davidson, P. Aubourg, D. A. Williams, Hematopoietic Stem-Cell Gene Therapy for Cerebral Adrenoleukodystrophy. N. Engl. J. Med. 377, 1630–1638 (2017).

61. A. Biffi, E. Montini, L. Lorioli, M. Cesani, F. Fumagalli, T. Plati, C. Baldoli, S. Martino, A. Calabria, S. Canale, F. Benedicenti, G. Vallanti, L. Biasco, S. Leo, N. Kabbara, G. Zanetti, W. B. Rizzo, N. A. L. Mehta, M. P. Cicalese, M. Casiraghi, J. J. Boelens, U. Del Carro, D. J. Dow, M. Schmidt, A. Assanelli, V. Neduva, C. Di Serio, E. Stupka, J. Gardner, C. von Kalle, C. Bordignon, F. Ciceri, A. Rovelli, M. G. Roncarolo, A. Aiuti, M. Sessa, L. Naldini, Lentiviral hematopoietic stem cell gene therapy benefits metachromatic leukodystrophy. Science 341, 1233158 (2013).

62. C. Yan, T. Zhao, H. Du, Lysosomal acid lipase in cancer. Oncoscience 2, 727–728 (2015).

63. K. Kuriwaki, H. Yoshida, Morphological characteristics of lipid accumulation in liver-constituting cells of acid lipase deficiency rats (Wolman’s disease model rats). Pathol. Int. 49, 291–297 (1999).

64. A. M. Davidoff, C. Y. C. Ng, J. Zhou, Y. Spence, A. C. Nathwani, Sex significantly influences transduction of murine liver by recombinant adeno-associated viral vectors through an androgen-dependent pathway. Blood 102, 480–488 (2003).

65. M. Piechnik, P. C. Amendum, K. Sawamoto, M. Stapleton, S. Khan, N. Fnu, V. Álvarez, A. M. H. Pachon, O. Danos, J. T. Bruder, S. Karumuthil-Melethil, S. Tomatsu, Sex Difference Leads to Differential Gene Expression Patterns and Therapeutic Efficacy in Mucopolysaccharidosis IVA Murine Model Receiving AAV8 Gene Therapy. Int. J. Mol. Sci. 23, 12693 (2022).

66. G. Ronzitti, G. Bortolussi, R. van Dijk, F. Collaud, S. Charles, C. Leborgne, P. Vidal, S. Martin, B. Gjata, M. S. Sola, L. van Wittenberghe, A. Vignaud, P. Veron, P. J. Bosma, A. F. Muro, F. Mingozzi, A translationally optimized AAV-UGT1A1 vector drives safe and long-lasting correction of Crigler-Najjar syndrome. Mol. Ther. Methods Clin. Dev. 3, 16049 (2016).

67. E. Ayuso, F. Mingozzi, F. Bosch, Production, purification and characterization of adeno-associated vectors. Curr. Gene Ther. 10, 423–436 (2010).

68. U.-P. Rohr, M.-A. Wulf, S. Stahn, U. Steidl, R. Haas, R. Kronenwett, Fast and reliable titration of recombinant adeno-associated virus type-2 using quantitative real-time PCR. J. Virol. Methods 106, 81–88 (2002).

69. P. Bankhead, M. B. Loughrey, J. A. Fernández, Y. Dombrowski, D. G. McArt, P. D. Dunne, S. McQuaid, R. T. Gray, L. J. Murray, H. G. Coleman, J. A. James, M. Salto-Tellez, P. W. Hamilton, QuPath: Open source software for digital pathology image analysis. Sci. Rep. 7, 16878 (2017).

70. R. Patro, G. Duggal, M. I. Love, R. A. Irizarry, C. Kingsford, Salmon provides fast and bias-aware quantification of transcript expression. Nat. Methods 14, 417–419 (2017).

71. M. I. Love, W. Huber, S. Anders, Moderated estimation of fold change and dispersion for RNA-seq data with DESeq2. Genome Biol. 15, 550 (2014).

72. T. Wu, E. Hu, S. Xu, M. Chen, P. Guo, Z. Dai, T. Feng, L. Zhou, W. Tang, L. Zhan, X. Fu, S. Liu, X. Bo, G. Yu, clusterProfiler 4.0: A universal enrichment tool for interpreting omics data. Innov. Camb. Mass 2, 100141 (2021).

73. G. Yu, Q.-Y. He, ReactomePA: an R/Bioconductor package for reactome pathway analysis and visualization. Mol. Biosyst. 12, 477–479 (2016).

74. C. Osto, I. Y. Benador, J. Ngo, M. Liesa, L. Stiles, R. Acin-Perez, O. S. Shirihai, Measuring Mitochondrial Respiration in Previously Frozen Biological Samples. Curr. Protoc. Cell Biol. 89, e116 (2020).

